# Development of Molecular Digital Twins Based on Ambient Ionization Mass Spectrometry Imaging for Application in Cancer Surgery

**DOI:** 10.1101/2024.12.08.627387

**Authors:** Yanis Zirem, Léa Ledoux, Nina Ogrinc, Laurine Lagache, Roland Bourette, Chann Lagadec, Paul Chaillou, Michel Salzet, Isabelle Fournier

## Abstract

Cancer surgery is a fundamental component of oncology treatment, its quality significantly impacts patient outcomes, influencing both relapse rates and survival. However, achieving this customization is contingent upon early collection of robust molecular data during surgery, providing accurate information for diagnosis, prognosis, and delineating surgical margins. The introduction of digital twin (DT) technology has recently opened a new era of precision and effectiveness in cancer surgery. Expanding from its successful implementations in the industrial sector, DT concept has evolved into a highly promising breakthrough in healthcare. Therefore, our study goal is on creating DT by using accurate and high-throughput molecular data obtained through mass spectrometry imaging. We developed a machine-learning-based pipeline that allow to depict infiltration of cancer cells into normal tissue that offer precise delineation of tumor margins thanks to SpiderMass. This process also enables the prediction of relative presence of bacterial strains in tumoral and healthy mammary glands.

## Introduction

Cancer surgery most often remains the first pillar of therapy in oncology. Surgery quality is of utmost importance because of the huge impact it has on cancer recurrence and patient survival. Besides, the therapeutic management choice will very often depend on the outcome of the surgery and the post-surgical assessment of excised tissues. The aim of any cancer surgery is to remove the cancer with an adequate margin of normal tissue but with minimal morbidity. During the procedure, surgeons cannot access in real-time to data that will help them discriminate normal tissues from tissues infiltrated with cancer cells. They thus must rely on their experience and training to make their decision. This lack of objective data lead, in application of the precautionary principle, to take wider surgical margins increasing morbidity and decreasing patient life quality. Conversely, surgical reintervention must have to be carried out for positive margins detected at the post-surgery diagnosis. Yet the gold standard procedure is based on the histopathological examination of the excised specimens. Intraoperative examination is possible and beneficial to improve the surgery, but due to the constraints of the process, it’s limited to a few parts of the tissues not well recapitulating the specimen globality. For logistic reasons and lack of pathologists, it is also very difficult to implement for all surgeries. The final diagnosis is therefore only made during the post-treatment examination and may lead to the discovery of positive margins or, conversely, the removal of tissue that was unnecessary. Besides, only excised tissues are examined and the status of the bedside or adjacent tissues remain unknown. Further improving the surgical oncology procedures implies moving forward more personalized surgery. However, tailoring the surgery is tightly linked to the ability to collect robust molecular data already by the time point of the procedure. Collecting accurate molecular information will help the surgeon by offering in real-time the knowledge on the tissue status that is central for the margin delineation or to find the loco-regional extension of the cancer (e.g. status of the sentinel node). Additionally, it could also be used to get a real-time diagnosis and prognosis which according to the type of solid tumor could be taken into consideration to adapt the surgery (e.g. detection of an aggressive cancer subtype).

Digital Twin (DT) represents a paradigm shift for precision medicine^1^ and paves the way in oncology to enter the era of precision cancer care^2^. If the concept of precision oncology is most often used for adapting the treatment of patients based on the administration of medicines, it is however an emerging concept in surgery. Introduced in 2010 by J. Wicks in the NASA roadmap^3^, the concept of DT initially found its roots in manufacturing and engineering, primarily serving purposes in product design, service management, and the real-time monitoring of industrial equipment^4^. Building on the successful applications within the industrial sector, the DT concept has now become one of the most promising advancements in healthcare with the creation of patients DT. When discussing digital twins in healthcare, the focus often revolves around the integration of connected devices like smart sensors to enhance data collection, like blood glucose levels, MRI, temperature, CT scans, cardiac electrophysiology or even clinical data^5^. This integration facilitates simulations with real-world scenarios, thereby reducing medical risks and costs, and enhancing the quality of diagnosis, treatment, and disease prediction. DT of patients help in developing a more personalized and precise medicine by gathering real-time monitoring and adaptation of treatments as well as improved management by predictions through machine learning. Cancer patients digital twins (CPDT) have proven to be of significant value in the field of oncology^6^. Their potential applications are therefore diverse, including monitoring, diagnosis, cancer surgery and development of therapeutics strategies tailored to individual patients^7^. Notably, there is a growing interest in exploring CPDT for the discovery of novel treatment targets and the prediction of drugs effects. Aggregating data from different modalities into a single representation provides indeed a more accurate overall view of patient characteristics, which can be used to predict response to treatment and improve care. On the other side, DT concept also find important application in surgery to better prepare the surgery, improve the post-surgical management using DT recorded during the surgery or using DT during the surgery as surgical^8–10^. For example, the anatomical structure of the patient can be augmented by molecular data to help the surgeon. In cancer surgery, this involves the creation of virtual replicas of a patient’s tumor and surrounding anatomical structures, using real-time data and advanced simulations. These digital counterparts enable oncologic surgeons to meticulously plan and execute procedures with an unprecedented level of precision, ultimately leading to enhanced patient outcomes^2,11^. The fusion of this digital duplicate with cutting-edge technologies like cloud computing, artificial intelligence (AI) and machine learning (ML) will facilitate the comprehensive understanding, analysis, manipulation of data and informed decision-making. For example, Angulo *et al.*^12^ have introduced a digital twin for monitoring lung cancer behavior in patients, tailoring healthcare interventions based on the disease’s specific impact on individuals. Bagaria *et al.*^13^ have proposed a DT for monitoring heart rate and galvanic response to prevent heart diseases. For instance, Meraghni *et al.*^14^ have suggested a DT for breast cancer (BC) using temperature sensor information collected by portable intelligent devices. In health treatments, DT can simulate new experimental decisions within a virtual “real” environment to assess treatment outcomes^15^. Using a digital twin combined with AI offers the chance to create personalized recommendations tailored to the specific circumstances of the patient^16,17^. Clinicians can then utilize these recommendations to make decisions that are not only more accurate but also highly personalized and effective.

Over the past decade mass spectrometry (MS) has emerged as an interesting technology to assist surgery. MS can be performed in-situ and provide non-targeted molecular data. As MS separates molecules according to their molecular weight, the MS spectra provide a profile of the molecules detected from the tissues based on hundreds of different compounds with various intensities. These molecular profiles have been shown to be very specific of the cell phenotypes. Hence the presence of cancer cells, or existence of different cancer subtypes driven by different pathophysiological mechanisms are traduced in the MS molecular profiles. Using Mass Spectrometry Imaging (MSI), these molecular profiles can be recorded in a systematic manner by scanning a region of interest (ROI) to get the spatial distribution of the molecules within this ROI. MSI has demonstrated to be an essential tool in cancer research, revolutionizing our ability to explore the molecular intricacies of tumor tissues^18,19^. By combining the precision of MS with spatial information, MSI allows researchers to create detailed maps of various biomolecules such as metabolites, lipids, and proteins within cancerous tissues^20,21^. This spatially resolved molecular data unveils the complex heterogeneity of tumors, shedding light on their metabolism, biomarker profiles, and microenvironment^22^. Importantly, ambient ionization mass spectrometry (AIMS) based technologies have emerged to enable in-man MS analysis during surgery^23–25^. Among AIMS technologies, only few were shown to be operable *in vivo* for surgical purposes^26^. Advantageously, SpiderMass technology^27^, which is promoting a micro-sampling of the tissue through resonant excitation of water molecules endogenous to the tissues, was demonstrated to be a micro-invasive, painless and contactless technology, for intraoperative analysis^27,28^. Thanks to the limited damage to the tissues (a few µm depth) and the contactless analysis, SpiderMass stands out as the only MS solution permitting in man MSI ^29,30^. MSI with the SpiderMass technology^30^ is obtained thanks to an integration of the laser probe of the system onto a 6-axis robotic arm which can be moved to scan tissues according to a defined raster. Interestingly, a distance sensor is also added into the head of the robotic arm to measure the sample topography as well. Thus, during a MSI acquisition, the system capture at the same time both the molecular data and the topography, providing access to the reconstruction of 3D molecular imaging maps.

Cancer surgery aims at removing the cancer but uniquely the diseased tissues while preserving normal ones. However, practitioners are often at a loss as they are not always able to determine the status of the tissue and make the best decision for the patient. Highly invasive surgery is associated with limited recurrence but significant surgical complications and high morbidity rates. Conversely, leaving cancerous tissues/cells after surgery is associated with much lower 5-year survival rates. Hence, the accurate delineation of the resection margins and the loco-regional extension of the disease are paramount to the quality of the surgery. The development of new technologies, such as AIMS, offers an interesting prospect for the surgeon in guiding his decisions. Nevertheless, due to its format and the large amount of information it contains, the MS data collected intraoperatively cannot be used directly and require an interpretation and have to be represented in such a way that the end-users can obtain the information they need in the most readable format.

This study thus focuses on a new concept of molecular digital twins for solid tumor surgery with the generation of digital twins from non-targeted molecular data based on MSI collected thanks to the SpiderMass technology. To create these CPDT, we combine the molecular data from MSI interpreted through a machine learning (ML)-based interpretation pipeline and their 3D virtual counterparts to offer a better delineation of the margins, and their infiltration by cancer cells, to the practician leading to a more tailored surgery. Classifying cells of homogeneous (or very closed) phenotypes has already been demonstrated for various intraoperative technologies including MS and can be achieved thanks to conventional ML algorithms^31–33^. However, interpreting mixed molecular profiles issued from mixed cell populations is above the capacity of simple classification and generally lead to false positive or negative results. More specifically, predicting the ratio of the different cells subpopulation in an image pixel require a more complex processing^34^. Hence, we are presenting the development of a robust and novel processing pipeline to generate these MS-based CPDT. With this one, we can predict the ratio of different cell population in each pixel of the molecular image, and for example the ratio of cancerous cells versus normal ones to get the margin delineation. Additionally, this pipeline allows also to predict the ratio of different bacterial strain in different region, and according to cell phenotypes, of the tissue. This can be applied to highlight the presence of specific bacterial populations according to cancer subtypes or in different areas of the tissues for diagnosis and prognosis purposes. To proof-of-concept these novel CPDT, we have worked with transgenic mice models, which spontaneously develop mammary tumors, as a mimic of triple negative breast cancer, and generated DT after scanning tumors from different mice. We are now investigating generating these scores in real-time and unrolling them on the topographic maps recorded during the MSI to reconstruct DT based on the MS Imaging scoring which can then be used by the physicians to personalize their surgery. We illuminate how molecular DT can reshape the landscape of cancer surgery, offering new avenues for enhanced positive patient outcomes and medical education.

## Results

### SpiderMass MS Imaging and molecular DT creation workflow

The CPDT based on MS Imaging data was proof-of-concept on transgenic TgC(1)3 mice showing spontaneous development of mammary gland tumors. The overall design and workflow of the generation of molecular DT is shown in **Figure 1**. The MS Imaging SpiderMass system used for the DT is presented in **Figure 1A-B** and corresponds to an upgrade from our first lab prototype^30^. Briefly, the handheld of a fibered OPO laser equipped with a 1m length optical fiber and emitting at 2.94 µm (to promote resonant excitation of water) is connected to the head of a robotic arm which also includes a distance sensor. A tygon tubing of 1.5m is also connected to the head of the robotic arm to aspire the aerosol generated during the desorption-ionization process upon laser irradiation and transport it back the to the MS instrument thanks to high vacuum inside the mass spectrometer. The MS instrument without its conventional ion source is equipped with an interface which process the aerosol and avoid extensive fouling of the instrument while helping with desolvating the aggregates contained within the aerosol. This enables MS spectra to be generated in real-time (ms scale). The robotic arm is programed to move above the sample surface, to perform point-to-point MSI acquisition while adjusting the focal distance. In the present version, we have improved the scanning system and data acquisition for simultaneously collecting both the topographical and the molecular data (**Figure 1C**). Additionally, a high-precision laser distance sensor with 5 µm accuracy in all dimensions was incorporated and the SpiderMass laser microprobe is placed at an angle of 30° to ensure the convergence of the two laser beams (distance sensor and MS laser) in the focal plane. The system is powered by a novel version of OPO midIR fibered laser based on an Opolette 2940 mid-IR pulsed laser (Opotek, USA, CA) which is designed for a plug & play connection through high energy SMA connectors of a metal jacketed fiber. The injection in the optical fiber is made in a N2 flowed chamber to prevent the presence of dust and hance extensive burning of the fiber entrance. On the other side, at the fiber outlet, a handheld is mounted which includes a focusing CaF2 lens with a 4 cm focal distance. For each experiment the post-mortem mice were place underneath the sensor and microprobe as shown in **Figure 1B**. Using this SpiderMass MSI prototype, we have created a complete workflow ranging from the data acquisition to the DT creation (**Figure 1D-H**). The data are automatically acquired using a homemade acquisition interface programed under MatLab in 20 to 30 min runs depending on the size (around one or more centimeters) of the region to be imaged (tumor and peritumoral area) (**Figure 1D**). Data can be collected either in the positive or negative ion mode, or both (one after each other) depending the cancer type and which mode provides the best molecular discrimination. These result in a topographical image and molecular data recorded in every pixel. Optical images of the samples are also collected prior to the acquisition. An example of collected optical and topographical images of tumors M15-T2 and M2-T1 are show in **Figure 1E**. The optical image shows two tumors from two different mice imaged with their corresponding topographies. Based on segmentation of molecular images plot in their corresponding topographical images, the molecular profiles of each class were used to train machine learning models (**Figure 1F-G**). **Figure 1F** shows an example of the segmentation obtained for the M3-T2 tumor, using Silhouette plot to find the number of clusters which best match the molecular data and in **Figure 1G** of a machine learning model, here shown in the for positive ion mode. The models can then be validated in blind from different samples and the molecular DT created by transforming the physical information into the virtual space using the topographical data. All the registered data are, therefore, combined and can be displayed according to various representations including Total Ion Current (TIC) map, molecular segmentation map, margin delineation map or even bacterioscoring maps, as exemplified for tumor M9-T1 in **Figure 1H**. While the training of the ML models from the initial sample cohort and their validation can take sometimes (typically a week weeks depending the size of the cohort and its availability), the creation of DTs by querying the models in real-time during an application of the technology is fast, with instant feedback to the user.

**Figure 1.**
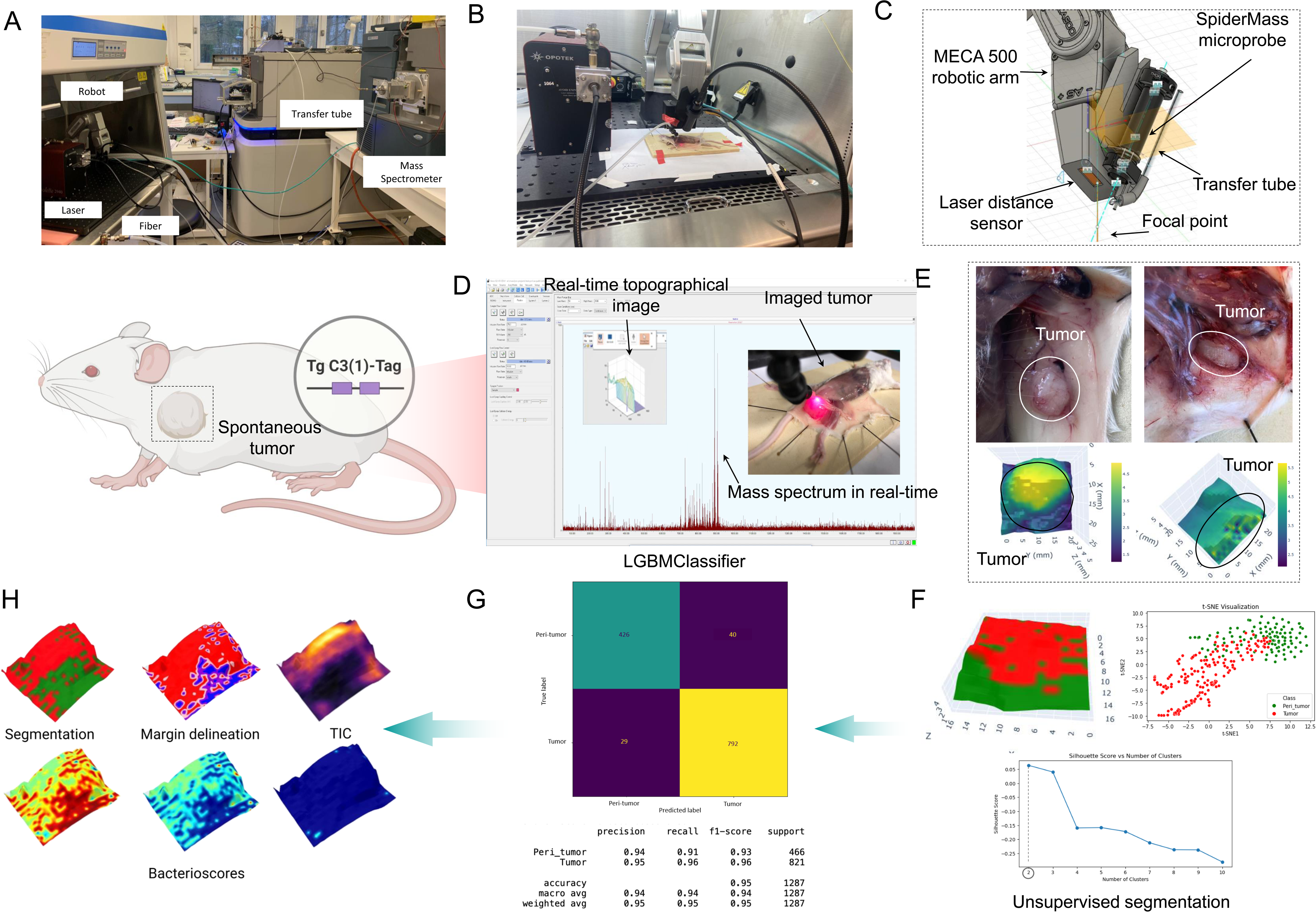
Workflow for the generation of the MS-based molecular digital twins. **(A-B)** Photo of the imaging setup including the Opolette 2940 laser with a reinforced jacketed fiber and an example of a mouse imaging experiment. The post-mortem mouse is exposed to reveal the tumor region and placed underneath the scanning system. **(C)** Schematic representation of the improved laser scanning system linked to the SpiderMass laser microprobe and transfer tubing on the robotic arm. **(D)** An example of 3D imaging acquisition. The image includes a real-time display of a real-time topography acquisition, mass spectrum and the photo of an imaged tumor. **(E)** Optical images of two mice with exposed tumor areas and corresponding topographical images obtained of the selected region. **(F)** Unsupervised segmentation to distinguish between tumoral and peritumoral areas. These clusters are then used to create the classification model. **(G)** Confusion matrix and classification report of the LGBMClassifier classification model built in positive ion mode from molecular profiles of tumoral and peri-tumoral regions. **(H)** Generation of different molecular digital twins based on various MS data.

### SpiderMass MS Imaging based DT training

A total of 12 transgenic mice were imaged post-mortem with SpiderMass after dissection. MS offers the possibility to work in both positive and negative ion modes, and some family of molecules can be better detected in one or the other mode. Most of experiments were conducted in negative ion mode involving 15 mammary glands from 9 animals while only 4 tumors from 3 mice were used for the positive ion mode. In addition, one healthy mammary gland from a separate healthy mouse was also subjected to MSI with SpiderMass, in negative ion mode only, as a negative control (**Table S1**). All data were collected at 500 µm spatial resolution. Photographs of one mouse tumor before and after an imaging experiment is shown in **Figure 2A**. Example of some optical images of mammary glands, with their corresponding topography images are presented **Figure 2B and Figure 3A** (framed in blue tumors used for training while in purple those for testing in blind). Three tumors were used for the training and one for the blind prediction in positive ion mode while in negative ion mode, fourteen were used for training purposes and two tumors were assigned to the validation blind set (**Table S1**). After mapping back the molecular data onto the 3D surfaces, image segmentation was performed by *k*-means ++ algorithm on all images and it highlights the presence of 2 clusters knowing that the choice of the optimal number of clusters was obtained using the Silhouette criterion (**Figure S1-S6**). These 2 clusters correspond to 2 molecularly distinct regions which correspond to tumor (red) and peritumoral region (green) (**Figure 2C and Figure 3B**). The assessment of clusters into peritumoral or tumoral tissue was visually conducted, leveraging the tumor’s topography—typically positioned higher contrasted with the peritumoral tissue located on lower sides. While this instance exemplified a straightforward class assignment, on intricate samples, multiple pathologists will assess each cluster prior creating the classification model. Furthermore, the distribution of the dataset for each tumor were depicted employing t-SNE dimension reduction. Two distinct point clouds are clearly observed, aligning with the presence of two clusters, one for the tumor and another for the peritumoral region (**Figure 3B**).

**Figure 2.**
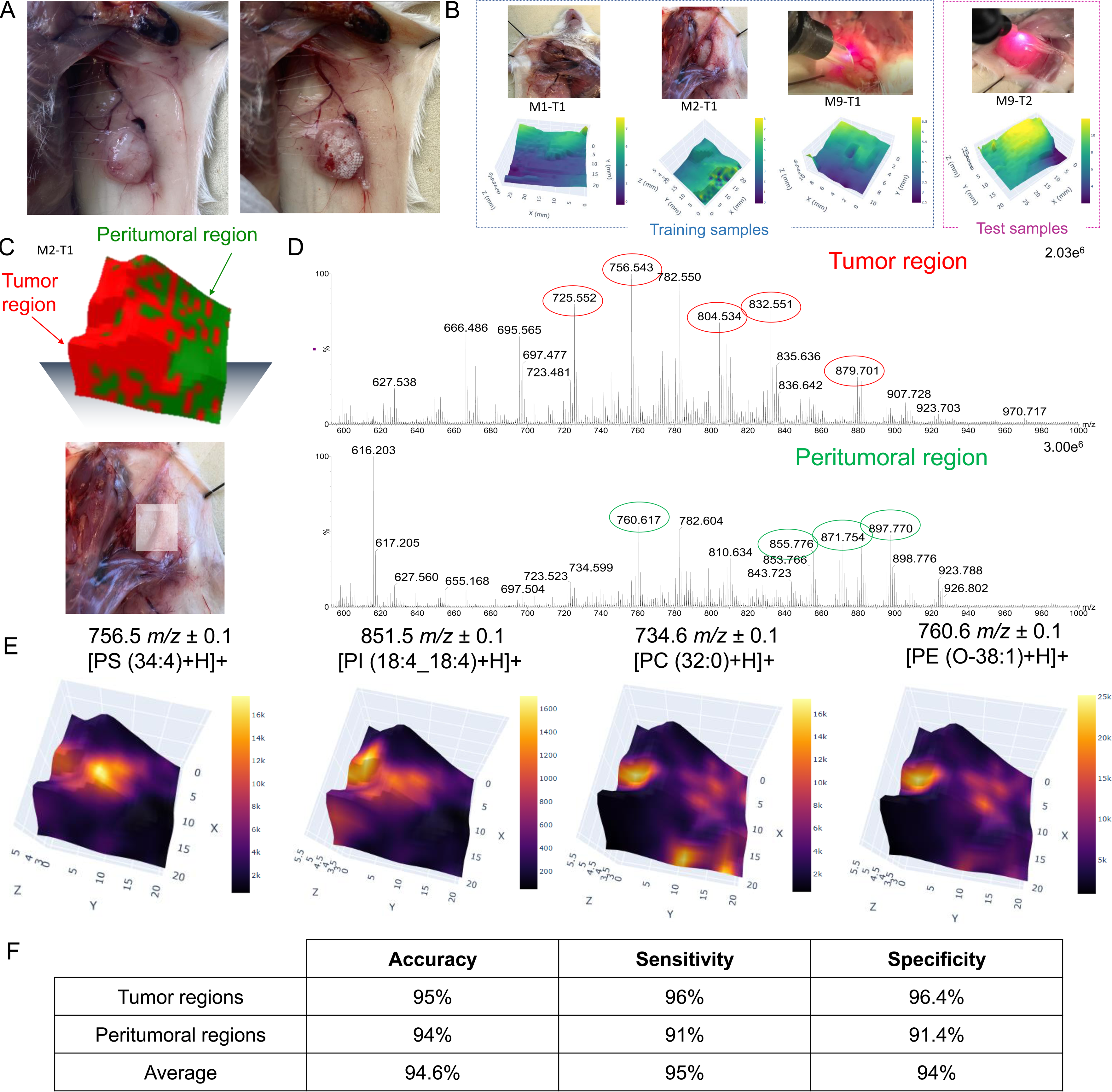
SpiderMass MS Imaging of TgC(1)3 mice mammary tumors in positive MS ion mode. **(A)** Photograph of the mouse tumor before and after the MSI experiment. The experiment leaves white dots of dehydration indicating the imaged area (tumor and subsequent peritumoral). **(B)** Several tumors from different mice were used as a training and validation cohort in positive ion mode. The optical images of tumors with the corresponding topography highlighted in the blue box served as training samples. The optical images and corresponding topography highlighted in the magenta box served as the validation cohort. **(C)** Imaged region of the M2-T1 tumor with the corresponding *k*-means ++ segmentation. This one depicts 2 clusters corresponding to tumor (red) and peritumoral region (green). **(D)** Extracted mass spectra from the tumor and peritumoral regions at 600-1000 *m/z*. The distinct peaks for each area are circled in green or red, respectively. **(E)** Single ion 3D reconstruction, on M2-T1 tumor, for *m/z* 756.5 ± 0.1, *m/z* 851.5 ± 0.1, *m/z* 734.6 ± 0.1 and *m/z* 760.6 ± 0.1. **(F)** Table with accuracies, sensitivities and specificities for tumor regions, peritumoral regions and in average after 20-fold cross-validation ending to 94.6% correct class prediction. Related to Table S1 and Figure S1.

**Figure 3.**
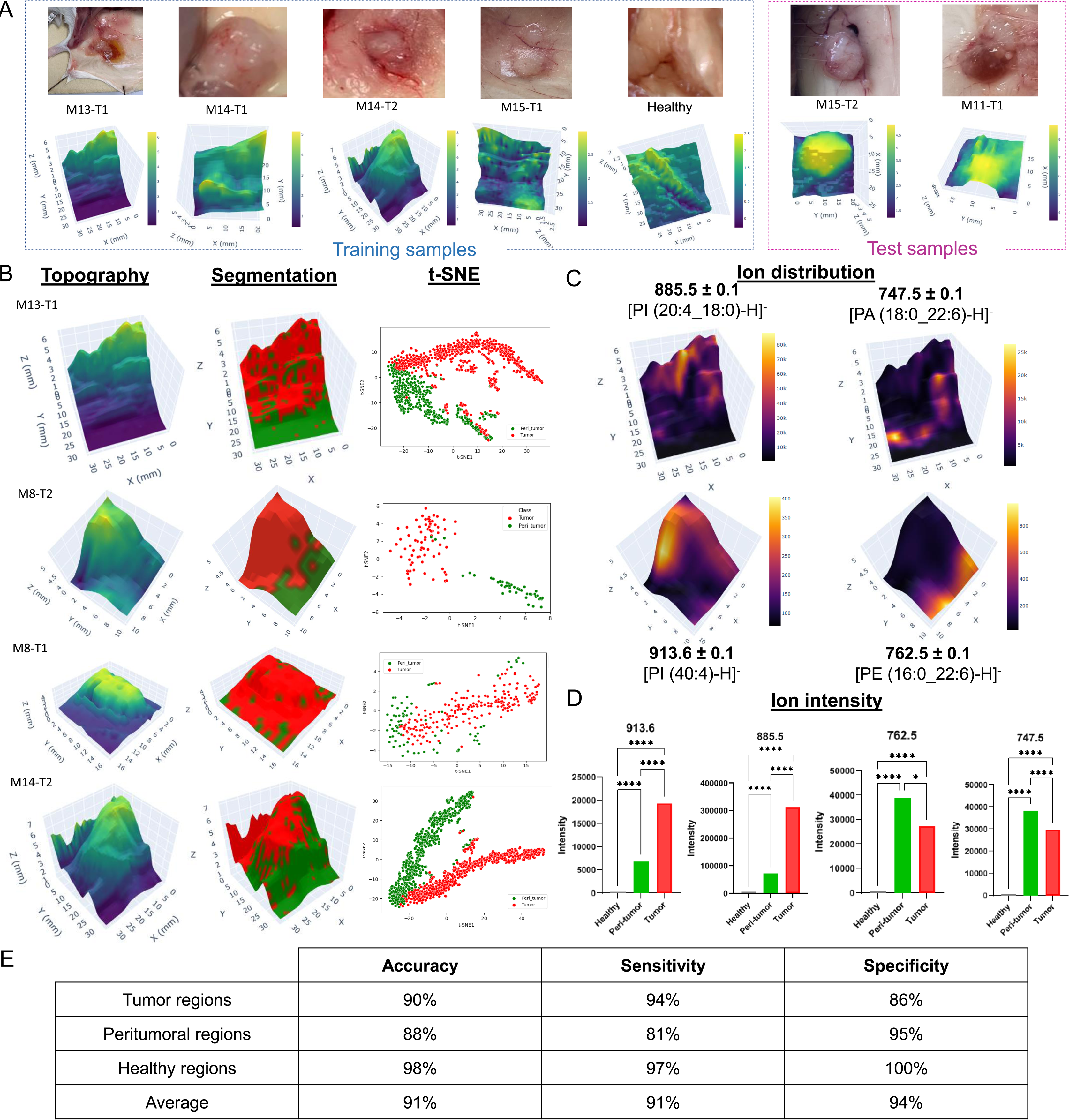
SpiderMass-MSI analysis on TgC(1)3 mice tumor regions in negative ion mode. **(A)** Several tumors from different mice were used as a training and validation cohort in negative ion mode. The optical images of tumors and a healthy mammary gland with the corresponding topography highlighted in the blue box served as training samples. The optical images and corresponding topography highlighted in the magenta box served as the validation cohort. **(B)** The 3D topographical image, the *k*-means ++ segmentation and the t-SNE visualization of 4 tumor examples (M13-T1, M8-T1, M8-T2 and M14-T2). Each individual segmentation reveals 2 distinct clusters mainly tumor (red) and peritumoral region (green). **(C)** The 3D selected ion images *m/z* 913.6 ± 0.1, *m/z* 885.5 ± 0.1, *m/z* 762.5 ± 0.1 and *m/z* 747.5 ± 0.1 on M13-T1 and M8-T2 respectively. **(D)** Corresponding boxplot representations of specific *m/z* values for each cluster. *p ≤ 0.05, **p ≤ 0.01, ***p ≤ 0.001, ****p ≤ 0.0001. **(E)** Table with accuracies, sensitivities and specificities for tumor regions, peritumoral regions, healthy regions and in average after 20-fold cross-validation ending to a 91% correct class prediction. Related to Table S1 and Figures S2-7.

Molecular differences between the two clusters are well observed in the extracted average mass spectra from the tumor and peritumoral region as shown for the *m/z* 600-1000 range in positive ion mode (**Figure 2D**). In a visual manner, some signals seem to be exclusive to one of the clusters, as the *m/z* 756.5 [PS (34:4)+H]^+^ and *m/z* 832.5 [PS (40:8)+H]^+^ which seems only present in the tumor regions in contrary to ions *m/z* 760.6 [PE (O-38:1)+H]^+^ and *m/z* 871.7 which seems specific of the peritumoral regions. More interestingly, some ions were found to be zone-exclusive using a Kruskall-Wallis significance test in both ion mode. In fact, in positive ion mode, as the **Figure 2E** show, the selected ion 3D image *m/z* 756.5 ± 0.1 and *m/z* 851.5 ± 0.1 [PI (18:4_18:4)+H]^+^ overlap with the tumor region while the ion image of *m/z* 734.6 ± 0.1 [PC (32:0)+H]^+^ and *m/z* 760.6 ± 0.1 overlay with the peritumoral region. In addition, in negative ion mode, the selected ion image of the *m/z* 913.6 ± 0.1 [PI (40:4)-H]^-^ and *m/z* 885.5 ± 0.1 [PI (20:4_18:0)-H]^-^ overlap with the tumor region, whereas the ion images of *m/z* 762.5 ± 0.1 [PE (16:0_22:6)-H]^-^ and *m/z* 747.5 ± 0.1 [PA (18:0_22:6)-H]^-^ overlay with the peritumoral region (**Figure 3C**). Furthermore, this region specificity is confirmed by the corresponding boxplot representations (**Figure 3D**). To be noted that the healthy mammary gland is defined by the absence of the ions which are found to be very abundant in either the tumor or peritumoral regions rather than being discriminated by the presence of specific ions.

Two classification models, in each ion mode, were then constructed using all corresponding MS spectra acquired from the training samples. LGBMClassifier was found to be the optimal algorithm for both ion mode datasets, according to our new developed pipeline^34^. Remarkably, the classification models obtained 95% and 91% of correct classification rate after 20-fold cross-validation, in positive and negative ion mode respectively. Moreover, the sensitivity of these classification models, based on unsupervised segmentation of molecular data, are 91% and 95% respectively in positive and negative ion mode while their specificities are 94% for both (**Figure 2F and Figure 3E**).

### SpiderMass based 3D DT from blind prediction

The built classification models were tested on additional tissues to the training cohort for the identification of tumoral versus peritumoral versus healthy regions in blind. The prediction scores resulting from the blind interrogation were plotted back onto the topographic maps to generate 3D prediction DT (**Figure 4**). In the positive MS mode (**Figure 4A**), blind prediction shows that the imaged mammary gland, M9-T1, is composed by 75.6% of tumor and 24.4% of peritumoral areas. These blind predictions from supervised ML approach were compared to unsupervised *k*-means ++ segmentation. The unsupervised approach gives 37.4% for peritumoral and 62.6 % for tumor area which is different from the supervised approach. The prediction similarity between supervised and non-supervised is then 64.2%. Similarly, in the negative MS ion mode (**Figure 4B**), two tumors were submitted to blind prediction. No normal tissues, as expected, are found in both mammary gland mice tumors. The first tumor, M11-T1, blind prediction reveals 71.9% tumoral and 27.9% peritumoral regions. A comparison with segmentation demonstrated a highly promising 90.03% similarity between the result for supervised versus unsupervised approaches. For the second tumor, M15-T2, the prediction indicated 78.6% tumoral and 21.3% peritumoral regions but 63.8% and 36.2% for the segmentation. This tumor was thus exhibiting only 58% similarity between supervised and non-supervised. This highlights that molecular DT based on the prediction of cell populations can be built either using supervised or non-supervised processing. The non-supervised approach is attractive because it does not require training, but on the other hand it doesn’t give any clue about the class correspondence. This demonstrates that employing molecular DT has a significant potential for oncological surgery, through a novel approach using real-time ambient mass spectrometry imaging.

**Figure 4.**
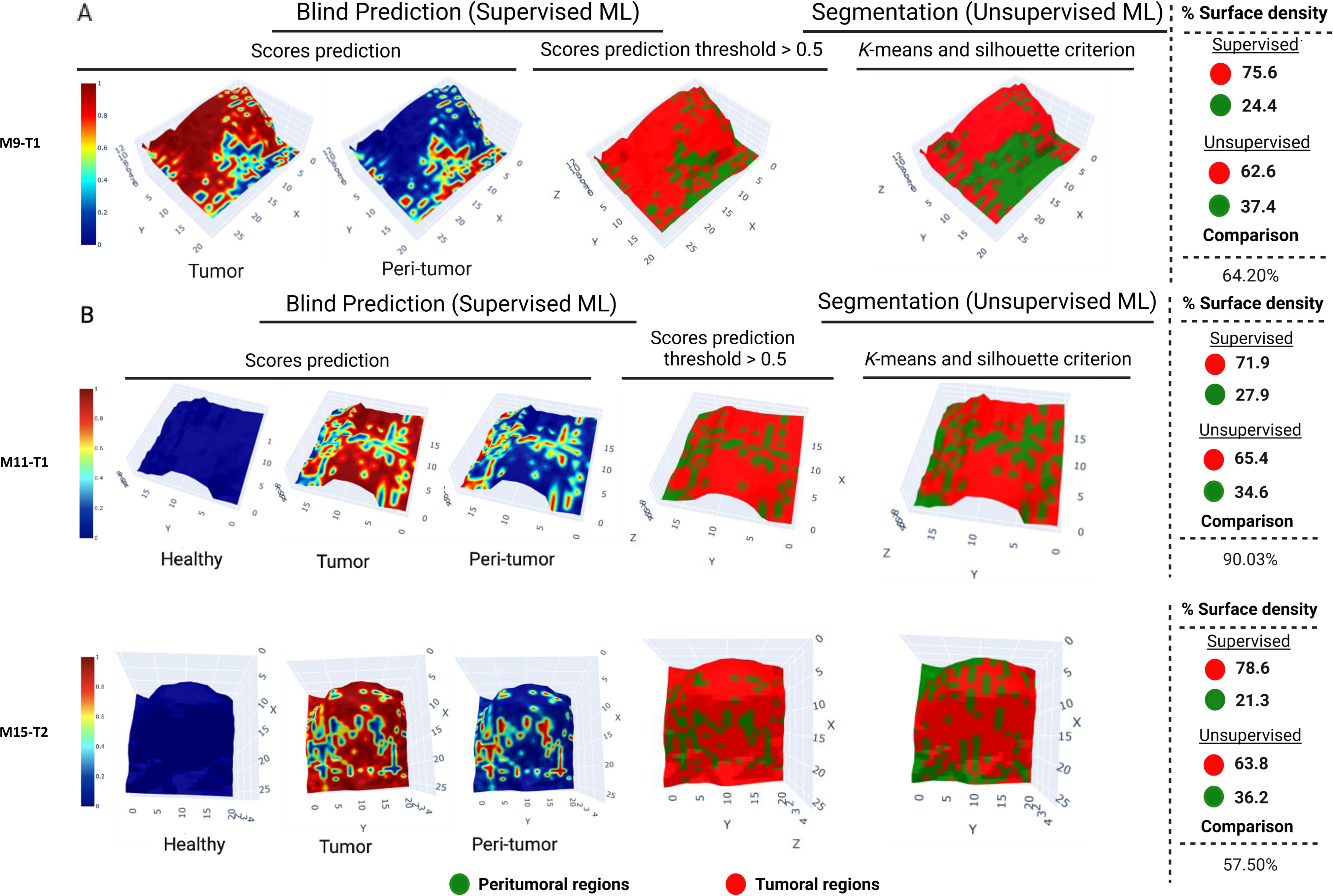
3D reconstruction of SpiderMass blind prediction. The prediction scores for tumor, peri-tumor, and healthy regions are reconstructed in 3D and the 3D map is also obtained with scores exceeding a threshold of 0.5, all this achieved through supervised machine learning. The corresponding segmentation obtained by unsupervised machine learning is also displayed. Furthermore, the surface density for each area is calculated and compared between the supervised and unsupervised approaches. **(A)** positive ion mode and **(B)** negative ion mode.

### Creating molecular DT for Margin delineation

Accurate margin delineation is the main objective of surgeon while removing the cancer tissues. Because of the delicate balance between the removal of cancerous tissue and the preservation of healthy tissue, the composition of normal and cancerous cells in the tissue must be precisely determined. Yet, SpiderMass technology analyzes at a spatial resolution of 500 µm which appears to be the best compromise between excision accuracy and time. However, with such a spatial resolution each analytical spot contains about 1000-2000 different cells which can be of different nature. This means that non-supervised segmentation will not be able to cope with predicting the cell population ratios within in a pixel of the SpiderMass image. To address this aspect, we have predicted the ratio of the different cell populations using the LGBMClassifier to construct the classification models. For margin delineation purposes, we only searched for the prediction of 2 cell populations, according to our model, which are cancer versus peritumoral cells. We exemplified the margin delineation DT by plotting these predicted ratios with different colors for 4 to 5 different ratio levels on the topographic maps, as illustrated in **Figure 5**. The four levels depict an intermediate region with tumor / peritumor ratios between 0.33 and 0.66, unlike the second model that portrays two in-between regions, each encompassing between 25% and 50% of either cancer or peritumoral cells. Each level ratio includes a range between 0 and 0.1, designated to emphasize regions deemed truly healthy, free from tumor infiltration. This processing was tested on a sample (M8-T2) from the training cohort (**Figure 5A**), showing the accuracy gained, compared to unsupervised segmentation only, for accurately finding the boundaries between peritumoral, tumoral and mixed cell area. More interestingly, the pipeline was also then applied on data issued from the three tumors employed for blind prediction (**Figure 5B**) in both positive and negative ion mode. In these three tissues (M9-T1, M11-T1 and M15-T2), locating the boundary between tumor and peritumoral tissue proves to be a delicate task without adequate molecular information. The DT based on predicted cell ratios showed that non-cancerous cells are infiltrating the tumor tissue. Only the M8-T2 and M9-T1 tumors contains areas with less than 10% tumor cells. This highlights the potential of DT based on predicted cell ratios to enhance precision enabling surgeons to make informed decisions regarding the optimal size for tumor resection, and ultimately enhancing patient prognosis and post-operative quality of life.

**Figure 5.**
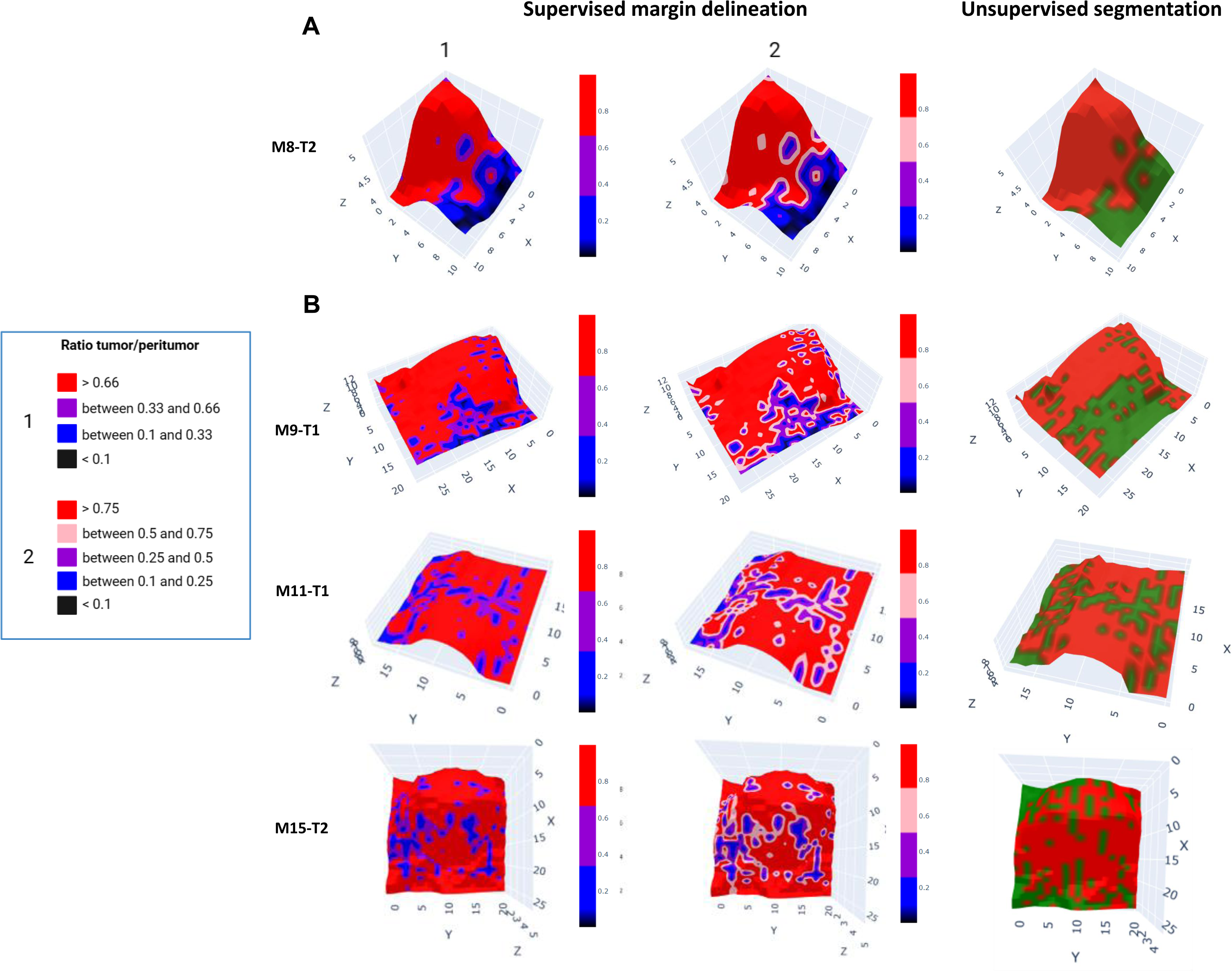
Margin delineation. The boundaries of tumor and peritumoral regions were delineated based on a ratio between tumor and peritumoral for **(A)** a mice tumor (M8-T2) used in the training of the classification model and for **(B)** three mice tumors (M9-T1, M11-T2 and M15-T2), not used in the classification model training, all using supervised blind prediction. For the color bar, the two options involve either a margin based on four ratio levels or on five ratio levels. Either illustrating an intermediate zone for ratios ranging from 0.33 to 0.66, or showing two intermediary regions, each covering between 25% and 50% of either peritumor or tumor areas.

### Bacterioscore-based DT

There are, over the past 5-years, growing evidence of the importance of the microbiota in cancer. Different bacterial populations are indeed found according to the cancer type and subtype^35^. It was also recently demonstrated that there is causal link between the presence of bacteria and the tumor behavior. Tissue microenvironment (TME) microbiota was shown to alter the biology of cell compartments and thus the immunity response as well as the migration of the cancer cells^36,37^. Microbiotic niches represent an interesting new avenue for therapy and for understanding treatment resistance. Therefore, we have been interested to evaluate the possible creation of DT based on the specific detection of bacterial strains within the tissues of the mouse mammary gland to provide a comprehensive assessment of the bacterial landscape using SpiderMass-MSI and develop a bacterioscore-based DT (**Figure 6**). To create the bacteriocore, we first analyzed by SpiderMass directly on agar plates three different bacterial strains (*S. infantis, S. lugdunensis* and *M. radiotolerans*) which were previously shown to be present in different breast cancers^38–40^. Using machine learning algorithms, we trained a model in both positive and negative MS ion mode using 70 MS spectra per bacterial strain, we achieved a correct classification rate of 100% in both training and cross-validation sets (**Figure 6A**). The training was then unrolled on the topographic molecular images of the tumor mammary glands. For each pixel of the image, it is possible to predict the ratio of the different bacteria within the pixel^34^. The obtained results furnish estimated scores for each bacterial strain, and ratio scores are calculated to determine the relative score presence of each bacterial type across the entire image and plot this result on the topographic map to create the bacterioscore-based DT. This innovative approach allowed the reconstruction of a 3D image for all tumors, providing the probability of presence of each bacterial strain expressed in each pixel. Indeed, more the pixel is red more the bacterium is present in the pixel (**Figure 6A**). Interestingly, the expression levels of bacteria were found to be associated with specific regions of the tissue and consistent results were obtained in both MS ion modes. All the predicted percentage in the different peritumoral, tumor and healthy regions for the different mammary glands imaged by SpiderMass can be found in **Table S2**. Specifically, the *S. infantis* strain exhibited a higher predicted abundance in tumor tissues (80%) compared to peritumoral (66%) and healthy tissues (4.3%) which is in line with previous results obtained by sequencing^35^. In contrast, *S. lugdunensis* and *M. radiotolerans* demonstrated higher expression in healthy tissues than in tumor tissues (**Figure 6B)**. There is a significant contrast in the expression levels of *M. radiotolerans*, as it shows a relative presence of 58% in a normal mammary gland, whereas its presence decreases to 10% in tissues adjacent to and within tumors. This correlates with finding from a study indicating that the bacterial strain *M. radiotolerans* is more commonly present in hormone-dependent (estrogen-positive) rather than in triple-negative breast cancer tissue, the transgenic mice models used in our study mimicking triple-negative subtype^41^.

**Figure 6.**
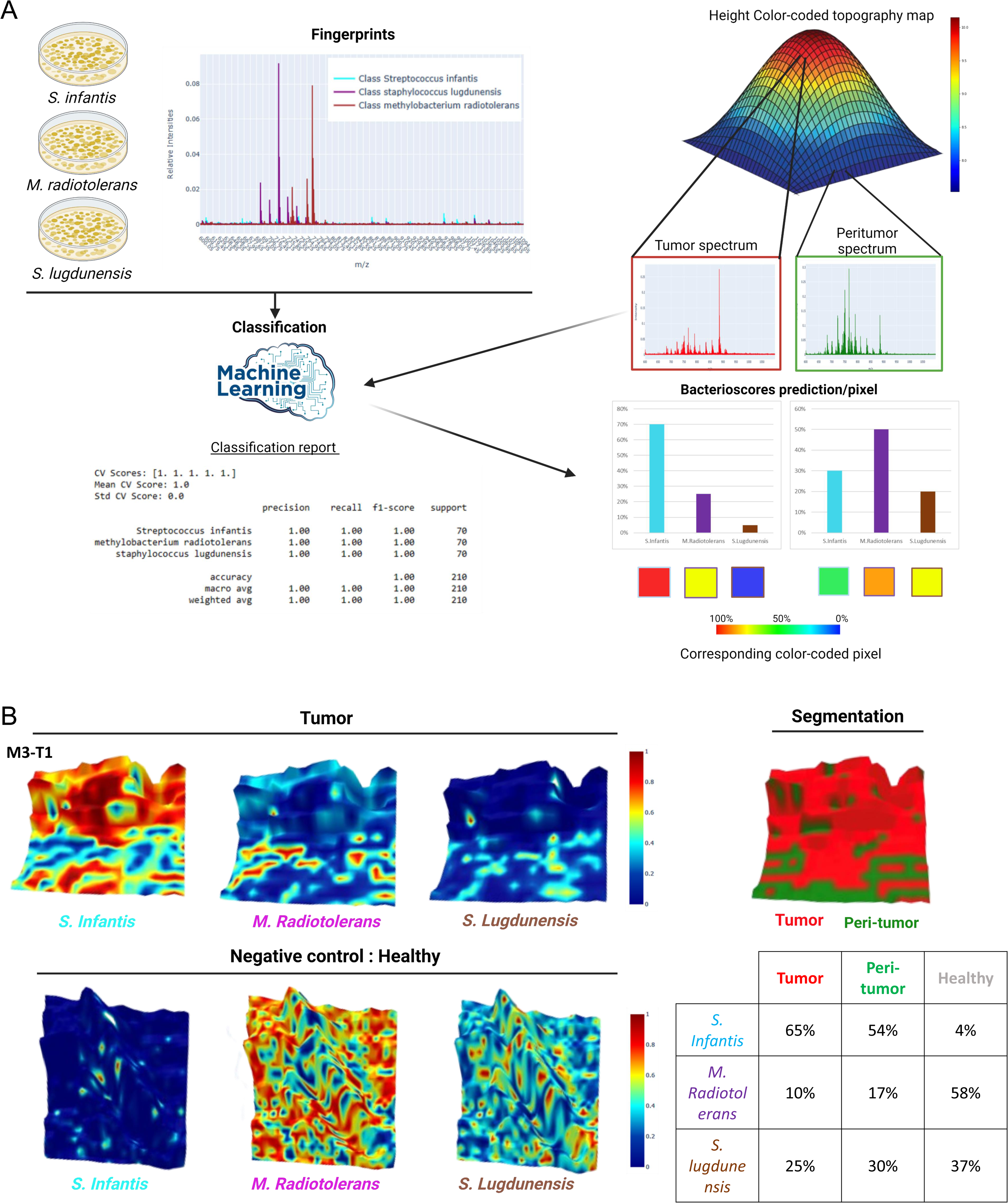
Automated bacterial strain recognition on 3D tumor. **(A)** Methodology employed to obtain the different bacterioscores prediction/pixel. **(B)** 3D reconstruction of bacterioscores for 3 bacterial strains in one mice tumor (M3-T1) and in one healthy mammary gland. Related to Table S2.

## Discussion

DT research is highly promising for customized patient care, with the potential to significantly impact the future landscape of healthcare innovation. A DT corresponds to a virtual embodiment of real-word objects or systems. In industry, DT were shown to be able to predict the behavior and response of a real-world object. In the case of cancers, evolution and response are linked to a complex interplay of different factors including the patient’s biology (i.e. genetics metabolism, etc.), lifestyle and external exposure factors (e.g. environment, pollution, etc.). Ideally, the creation of DT should consider all of these parameters to provide accurate prediction of the patient’s evolution with cancer and their response to treatment. It is extremely difficult though to obtain all this data on patients including their lifestyle and the environmental pressures to which they were subjected over the decades preceding the development of the cancer. However, these external factors influence the human biology and are reflected in the signaling pathways and levels of different biomolecules and xenobiotics which are expressed in patients’ tissues and fluids. The impact of external factors on the internal biology of patients demonstrates the value of using molecular data (derived from omics) to predict the evolution of the organism in relation to pathophysiological mechanisms. The development of CPDTs based on omics data is therefore of particular interest in cancer management by accurate forecasting. In the field of precision medicine, this is generally foreseen for adapting patient treatment without including the initial surgery within the therapy pipeline. Nonetheless, in solid tumors, the quality of the surgery is known to greatly impact patient prognosis and accurate diagnosis and prognosis by the time of surgery, opens the door a better tailored surgery as well as improved management. Thus, we have been investigating the development of CPDT based on lipidomic data issued from MS as potential tool for advanced diagnosis and prognosis which could be presented to guide surgeons in optimizing their procedure and patient management. Thanks to a recently developed ambient-ionization MS device the SpiderMass, lipidomic data can be monitored in real-time in-man. These data are obtained through the coupling to a robotic arm, in the imaging mode by scanning the laser probe of the SpiderMass above the ROI with the arm. From these molecular imaging data, we can create CPDT thanks to machine learning and present them as 3D map which are interpreted for the clinician. In the present study we have demonstrated the creation of these CPDT for surgery by analyzing TgC(1)3 mice which spontaneously develop mammary gland tumors and mimic triple negative BC. The mammary glands with cancer present two main regions which are the tumor area versus the peritumoral tissues. We show that a clear discrimination between the tumoral and peritumoral region is possible thanks to segmentation and involve ions which are different between the 2 regions and distinct from those found in the mammary gland of a normal mouse which is not a transgenic model. The silhouette scores and tests confirm that only 2 clusters are expected from the image data as awaited for these model tissues. Similar results are obtained in both positive and negative MS ion mode; discriminative lipids being different in the 2 modes though. Thus, these data can be used to train a classification model which enable to classify the tumoral and the peritumoral area versus the normal tissues, reaching up to respectively 95% and 91% in positive and negative MS ion modes. Interestingly, the model reaches a sensitivity of 95% and a specificity of 94% in positive ion mode while 91% sensitivity and 94% specificity are obtained in negative ion mode. These models are also validated in blind with class recognition and >90% similarity is obtained between the blind prediction and the non-supervised segmentation. *Ex vivo* part of the study can take several weeks to be done (to obtain the optimal classification model), yet querying the model in real time *in vivo* requires only a few milliseconds. The deployment of the prediction map in 3D leads to an easy-to-read representation that could be project in the surgery room for surgery guidance. However, affecting each image pixel to a single class is not sufficiently accurate for margin delineation which is one of the main objectives of the cancer surgeon. Indeed, each pixel which corresponds to a laser spot size of 500 µm, which appears to be the best compromise between excision accuracy and time for the end-user, includes 1000-2000 cells which can differ in their nature and phenotypes. To address this problem, we have developed an approach to predict the cell ratio of each pixel, here applied to 2 cell type situations with the tumor versus peritumoral cells. Thanks to this approach, we can offer CPDT presenting the area with different cell ratios highlighting area with only tumor cells versus area with only peritumoral cells as well as intermediate margin areas with different ratios of cell populations. The DT demonstrates that the peritumoral regions still contain cancer cells and that this tissue, despite presenting a different molecular profile and being segmented separately, cannot be consider as normal and should better be removed during the surgical process. It demonstrates that these CPDTs based on the ratio of cells map in 3D provide the surgeons with the right information to define the excision margins. Since images during surgery need to be acquire in a limited time (5-10 min maximum) instead of acquiring high spatial resolution (e.g. single cell) molecular images which would take far too much time (several hours), we have opted for a different approach where we keep lower spatial resolution (a few hundred micrometers) but we predict the ratio of the different cell populations within these pixels which contains a few thousands of cells including cancer and normal cells as well as immune cell infiltrated populations. To validate this approach, we have completed our study by some experiments on ovarian cancer tissues performed in 2D from which we have cross-validated the results from the prediction based on the SpiderMass MSI by a targeted approach using MALDI-MSI IHC (**Figure S8**). With the MALDI-IHC we can image in multiplex the distribution of several specific markers which target the different cell populations (cancer cells, conjunctive tissues and immune cells) and validate the predicted distribution for these cells from the SpiderMass molecular MSI data. Lastly, there are increased evidence of the role of bacteria in the onset of BC and its evolution. We wanted to assess the sensitivity of the SpiderMass technology to assess the presence and distribution of bacterial population in cancer tissues. By analyzing grown population of bacteria known to be present in BC, we were able to predict their presence in the molecular profiles recorded from the mammary gland tissues of transgenic and normal mice. By scoring their presence in each pixel, it was possible to create CPDT based on 3D maps bacterioscoring. Interestingly, these maps reveal that the *S. infantis* bacterial strain, which is known from previous work to be one of the most abundant bacteria found in BC^35^, is highly present in the cancer region with a limited presence in peritumoral tissues and almost completely absent from mammary glands of normal non-transgenic mice. Conversely, *M. radiotolerans* and *S. lugdunensis* are found in peritumoral regions with limited presence even for *S. lugdunensis*, while there are highly abundant in normal mammary glands. These 2 last bacteria were also described in cancer in human tissues; however here we used mice for the study which might explain the difference of the bacteria expression. In addition, other study indicating that *M. radiotolerans* was more prevalent in estrogen-positive BC tissue compared to triple-negative BC tissue^41^. Certain bacteria are shown to be associated with BC subtypes and immune cell niches in the tissue and could relate to the aggressivity of the cancer. In this context, knowing the cancer aggressivity would be an important information for the surgeon to consider tailoring the surgery within mind the future evolution of the patient. We have made yet the demonstration from transgenic mice model and we need in the next step to look forward to translate these developments to in-man application at the surgery room. For these, we will need to take further steps to enable the DT resulting from the model interpretation in real-time. Additionally, we need to address the speed of image acquisition to increase from the current 2.6 pixel/s screened. This will be made possible by continuous rather than spot to spot acquisition. Lastly, the bacterioscoring is extremely interesting and promising though we need to dig more into the microbiota of human BC by screening for more bacterial strains to better determine the location of different bacteria population and better understand their role in the cancer onset and progression. Finally, we demonstrate the interest of developing CPDT based on MS imaging data processed by machine learning and unrolled as 3D maps. Several CPDT representations will be accessible to the surgeon who will only have to pick the one thatbest match the needs (margin delineation, diagnosis, prognosis…) according to the cancer type, subtypeand the nature of the surgery. In the future, we are looking forward to decreasing the acquisition time using continuous acquisition mode to enable molecular topographic MSI data to be recorded in no more than 5 min followed by instant visualization of the DT. Hence, we do believe that these MSI-basedCPDT will be very useful in the future for cancer surgery guidance giving an accurate hand to the surgeon and promoting the evolution towards precision surgery.

## Acknowledgments

This research was supported by grants from Ministère de l’Enseignement Supérieur et de la Recherche (MESR), Inserm specific funding for SpiderMass project (I.F.), Inserm and Institut Universitaire de France (I.F.), Agence Nationale de la Recherche ANR-22-CE29-0016-01, DEADPOOL (I.F.). N.O. post-doctoral fellowship was funded by University of Lille Excellence Initiative from ERC Generator. L.L. post-doctoral fellowship is funded by Inserm from the Messidore SITH project. This project was supported by the Contrat de Plan Etat-Région 2021-2027 of the Hauts-de-France region in the frame of the CPER TechSanté (IF) and the region Council Hauts de France Multi-omics grant (L.La.).

## Author contributions

L.L., Y.Z., N.O. and I.F. wrote the manuscript original draft. I.F. and M.S. designed the experiments. N.O., L.L. and L.La. performed the experiments. R.B. is promotor of the mice model. C.L. collected the mice and performed the dissection. P.C. developed the imaging interface. Y.Z. designed the data processing. Y.Z. and L.L. analyzed the data. I.F., M.S., Y.Z., and L.L. and corrected the manuscript. I.F. promoted and supervised the project. I.F. and M.S. provided the funding.

## MATERIAL and METHODS

### KEY RESOURCE TABLE

#### Key resources table

| REAGENT or RESOURCE | SOURCE | IDENTIFIER |
| --- | --- | --- |
| Bacterial and virus strains |  |  |
| *Methylobacterium radiotolerans* | ATCC | ATCC 27329 |
| *Staphylococcus lugdunensis* | ATCC | ATCC 49576 |
| *Streptococcus infantis* | ATCC | ATCC 700779 |
| Chemicals, peptides, and recombinant proteins |  |  |
| Tryptic soy agar | BD | Cat#236950 |
| Nutrient agar | BD | Cat#213000 |
| Deposited data |  |  |
| The raw data from SpiderMass analysis has been deposited to the Harvard dataverse site. | This paper | Harvard dataverse : https://doi.org/10.7910/DVN/ZMUZ61 |
| Experimental models: Organisms/strains |  |  |
| FVB-Tg(C3-1-Tag)cJeg/JegJ mice | Charles River Laboratories (L'Arbresle, France) | / |
| One healthy female mouse | Animal Facility of University of Lille | / |
| Software and algorithms |  |  |
| Python 3.11.5 | Open source | RRID:SCR_008394<br>https://www.python.org |
| Jupyter Notebook 6.5.4 | Open source | https://jupyter.org |
| GraphPad Prism v9.5.1 | GraphPad software | RRID:SCR_000306<br>https://www.graphpad.com |
| Matlab R2019a | MathWorks | RRID:SCR_001622<br>https://matlab.mathworks.com |
| MassLynx 4.1 | Waters | RRID:SCR_014271<br>https://www.waters.com/waters/fr_FR/Logiciel-de-Spectrometrie-de-Masse-MassLynx/nav.htm?cid=513662&locale=fr_FR |
| The original code has been deposited on GitHub. The script for MSI SpiderMass dataset analysis was released and archived in Zenodo | This paper | https://github.com/yanisZirem/3D-MS-Imaging_data-analysis.git |
| The original code is available on GitHub and archived in Zenodo. This code was used to create the classification model based on MSI dataset. | Zirem *et al.* Cell reports medicine. 2024<sup>1</sup> | https://zenodo.org/doi/10.5281/zenodo.10656851 |
| The original code has been deposited on GitHub. The script for bacterioscore-based pipeline was released and archived in Zenodo | This paper | https://github.com/yanisZirem/Bacterioscores.git |

### RESOURCE AVAIBILITY

#### Lead Contact

Further information and requests for resources and reagents should be directed to and will be fulfilled by the lead contact, Isabelle Fournier.

#### Materials availability

This study did not generate new unique reagents.

#### Data and code availability

All SpiderMass raw/imzmL data (bacteria MS spectra and MS imaging data) and corresponding map files for each MSI tumor imaged have been deposited at Harvard dataverse and are publicly available as of the date of publication. DOI is listed in the key resources table.

All original code (3D MSI data analysis and Bacterioscore) has been deposited at GitHub (https://github.com/yanisZirem/3D-MS-Imaging_data-analysis.git and https://github.com/yanisZirem/Bacterioscores.git) and will be archived in Zenodo. If you have any questions or feedback, please contact.

Any additional information required to reanalyse the data reported in this paper is available from the lead contact upon request.

### EXPERIMENTAL MODEL AND SUBJECT DETAILS

#### TgC(1)3 mice model

Twelve FVB-Tg(C3-1-Tag)cJeg/JegJ mice were obtained from Charles River Laboratories (L’Arbresle, France). They were housed in the Pasteur Institute animal facility (PLETHA) and bred under a protocol approved by the Animal Protocol Review Committees of the Pasteur Institute (Lille, France) following European regulation (#25871-20200522117321730). For genotyping, ear clips were digested in lysis buffer (KAPA Biosystems). Samples were employed for PCR reactions to amplify T antigen cDNA using the primers TA1: 5’-GACCTGTGGCTGAGTTTGCTCA-3’ and TA2: 5’-GCTTTATTTGTAACCATTATAAG-3’. Products of the amplification were analyzed by agarose gel electrophoresis. As previously described, female mice expressing T antigen developed multiple mammary tumors by 4–5 months of age^42^. One healthy female mouse was obtained from the Animal facility of University of Lille. They were euthanized by exposure to a rising concentration of CO_2_.

#### Bacterial strain

Several bacterial strains (*Streptococcus infantis*, *Staphylococcus lugdunensis*, *Methylobacterium radiotolerans*) were obtained from the American Type Culture Collection (ATCC).

### METHODS DETAILS

#### TgC(1)3 mice model analysis

All of the mice were used for post-mortem imaging right after being sacrificed. Indeed, the mice were attached to the polystyrene foam using needles. The skin was notched with scissors without opening the peritoneum. To guide the scissors, a Brodie fistula director grooved with probe end was introduced between the skin and the peritoneum. After opening the shin, it was attached to the polystyrene foam using needles, thus exposing the mammary glands for 3D imaging analysis.

#### SpiderMass MS Imaging

The overall layout of the instrument setup has already been covered elsewhere^43^. In addition, here, the laser system used was an Opolette 2940 laser with a reinforced jacketed fiber. To perform imaging analysis, the Spider-Mass microprobe was coupled to a commercially available stiff 6D-axis precision MECA robotic arm (MECADEMIC, Montreal, Canada) described in a previous work^30^. The spatial step size was set either to 500 or 250 µm by achieving oversampling. The mass-range was fixed between *m/z* 100-1500. The acquisition sequence was composed of 3 consecutive laser shots and 3 seconds between each step. The laser bursts and the spectrometer acquisition were automatically triggered through a MATLAB in-house user interface developed for the robotic WALDI-MSI^30^. The data was acquired in positive and negative, sensitivity ion mode on a Xevo (G2-S, Q-TOF, Waters, Manchester, UK) mass analyzer through the REIMS prototype interface with biocompatible 2 meters Tygon tubing.

#### MS/MS analysis

SpiderMass technology facilitated the MS/MS investigation using the Xevo G2-S instrument from Waters. MS/MS spectra were recorded following the isolation of the precursor ion, which were then subjected to collision-induced dissociation (CID) in the transfer cell. The collision energy used for this ranged from 30 to 40 eV, depending on the specific precursor ion chosen. To identify lipids, a manual annotation process was employed, guided by fragmentation spectra guidelines, and the results were compared against databases such as LipidMaps, Alex123, MetFrag database^44^, and relevant literature.

#### Bacterial strain analysis

The bacterial strain used in this study are Streptococcus infantis (ATCC 700779), *Staphylococcus lugdunensis* (ATCC 49576), *Methylobacterium radiotolerans* (ATCC 27329). *S. infantis* and *S. lugdunensis* were cultured on Tryptic soy (BD 236950) agar plates when *M. radiotolerans* was cultured on Nutrient (BD 213000) agar plates, at 37°C for 24 hours. The SpiderMass analysis was made directly in the Petri dish in which the bacteria were grown. Several colonies were analyzed to get a good representativeness. To enable a good match of the molecular profiles, the SpiderMass MS spectra of the grown bacteria were obtained from the same spot size as the tissues.

#### Data processing and analysis

##### Image processing

The raw data files generated by the Spidermass instrument in both ionization modes were initially converted into the imzML format^19^ using MSConvert (Proteowizard ToolKit)^45^. To conduct data analysis, all imzML data files were imported into Python using Pyimzl library and pyimzml.ImzMLParser module. Once the data was imported, several pre-processing steps were applied to the spectra. These preprocessing steps involved normalization of total ion count (TIC) and binning to a window of 0.1 *m/z*. All final data sets contain 5000 *m/z* data points with the corresponding surface using x and y pixels coordinates. Indeed, for every raw file, the relevant map file was retrieved to extract topographic details (z-values) from the 3D image in CSV format. Ultimately, a 3D map was created using the provided z-values, accompanied by an interactive surface plot generated using the Plotly library. The resulting 3D plot facilitated the visualization of MSI data, including the exploration of spatial distributions of ions, TIC map, segmentation map, prediction label map as well as bacterioscoring map.

##### Segmentation

Spatial segmentation of SpiderMass imaging data was performed individually using the *k*-means++ algorithm from the scikit-learn library. This comprehensive spatial segmentation enabled the identification of regions of interest. To estimate the right number of clusters, not subjectively, the Silhouette criterion was employed. This criterion served multiple purposes, aiding in finding the optimal number of clusters and assessing their stability and compactness. Spectra associated with clusters, representing healthy, tumoral, or peritumoral regions, were labeled and grouped in a CSV file, forming the basis for building the classification model in both ionization modes.

##### Classification model and blind prediction^34^

The Lazy Predict library (https://lazypredict.readthedocs.io/en/latest/) was used to build multiple models from the scikit-learn library, encompassing 24 classifiers including linear models, ensembles, gradient boosting and neuronal network. The LGBMClassifier (Light Gradient Boosting Machine)^46^ emerged as the optimal model. Model assessment involved a 20% out, a 5-fold cross-validation and a 20-fold cross-validation using the KFold and cross_val_score functions. The classification report was generated, and the ConfusionMatrixDisplay function from the matplotlib library was used to visualized the confusion matrix. The optimal model was stored and blind predictions were executed using the joblib library’s dump and load functions.

##### Margin delineation

The constructed models were employed to predict the class of each pixel in an image using the “predict” function, which returns the class label with an argmax greater than 0.5. Consequently, pixels belonging to the same class are assigned the same color. To delineate the margin between classes, such as between a tumor and its surrounding peri-tumor region, the probability scores returned by the predict_proba function are utilized. The LGBM model effectively handles the predict_proba function. For models that do not inherently support probabilities, the CalibratedClassifierCV method could be used to make them probabilistic. The predict_proba method return continuous values (or predicted probabilities) that represent the likelihood of each new input belonging to each class (normal vs cancer). This is based on the features (ions) more or less present in the new spectrum compared to the characteristic spectrum of each class (tumor and healthy).

##### Bacterioscore

The bacterioscoring model underwent training using either the SGD Python library in negative ion mode or Ridge Classifier (sigmoid method) in positive ion mode. Both model used cell spectra within the *m/z* range of 600 to 1100 in both ion modes. The spectra were cathegorized into distinct bacterial strains, including *Streptococcus infantis*, *Staphylococcus lugdunensis*, and *Methylobacterium radiotolerans*, each consisting of 70 spectra. To predict the bacterial strain in SpiderMass images, the model’s predict_proba function was employed, offering probability estimates for bacterial strains and facilitating a nuanced understanding of the likelihood of each strain’s presence in the local environment. Moreover, ratio scores were calculated to estimate the relative presence of each bacterial strain across the entire image. These scores were determined by summing the scores for each strain and dividing it by the sum of the total scores across all labels. The ratios provided insights into the distribution of the trained bacterial strains throughout the image, offering a comprehensive assessment of the bacterial landscape in the analyzed sample.

##### Statistical tests

The discriminative *m/z* of the different segmentation’s clusters were found using a non-parametric statistical test Kruskal-Wallis with Bonferroni correction, employing the stats.kruskal function from the scipy library. First, a peak picking algorithm, the find_peaks_cwt function, was applied to remove instrument noise using a signal/noise ratio > 10. The corresponding box plots were then generated using GraphPad software where the legend correspond to *p value ≤ 0.05, **p value ≤\ 0.01, ***p value ≤ 0.001, ****p value ≤ 0.0001.

### QUANTIFICATION AND STATISTICAL ANALYSIS

Three datasets were employed for training, cross-validation, and testing of all classification models, including those for tissue type, and bacterioscore. Evaluation involved a 20% validation split and 20-fold cross-validation, with a classification report providing metrics such as accuracy, recall, precision, and F1 score. To assess the statistical significance of biomarkers, a non-parametric Kruskal-Wallis test was employed. Bonferroni corrections were applied to adjust p-values for multiple comparisons. Values are presented as medians and visualized through box plots.

## Supplemental figures legends

**Table S1.** Overview of the mice tumor cohort for the training and the testing of classification model in both ion modes. Related to Figures 2 and 3.

**Figure S1.** (A) Topographic map, (B) TIC map, (C) segmentation map, (D) silhouette score, (E) silhouette plot, (F) t-SNE and (G) average spectra of each clusters for 3 mice tumors (M1-T1, M2-T1 and M9-T1) used for the training in positive ion mode. Related to Figure 2.

**Figure S2.** (A) Topographic map, (B) TIC map, (C) segmentation map, (D) silhouette score, (E) silhouette plot, (F) t-SNE and (G) average spectra of each clusters 3 mice tumors (M3-T1, M3-T2 and M3-T3) used for the training in negative ion mode. Related to Figure 3.

**Figure S3.** (A) Topographic map, (B) TIC map, (C) segmentation map, (D) silhouette score, (E) silhouette plot, (F) t-SNE and (G) average spectra of each clusters 3 mice tumors (M4-T1, M7-T1 and M7-T2) used for the training in negative ion mode. Related to Figure 3.

**Figure S4.** (A) Topographic map, (B) TIC map, (C) segmentation map, (D) silhouette score, (E) silhouette plot, (F) t-SNE and (G) average spectra of each clusters 3 mice tumors (M8-T1, M8-T2 and M12-T1) used for the training in negative ion mode. Related to Figure 3.

**Figure S5.** (A) Topographic map, (B) TIC map, (C) segmentation map, (D) silhouette score, (E) silhouette plot, (F) t-SNE and (G) average spectra of each clusters 3 mice tumors (M13-T1, M14-T1 and M14-T2) used for the training in negative ion mode. Related to Figure 3.

**Figure S6.** (A) Topographic map, (B) TIC map, (C) segmentation map, (D) silhouette score, (E) silhouette plot, (F) t-SNE and (G) average spectra of each clusters 1 mice tumor (M15-T1) used for the training in negative ion mode. Related to Figure 3.

**Figure S7.** (A) Topographic map, (B) TIC map, (C) segmentation map, (D) silhouette score, (E) silhouette plot, (F) t-SNE and (G) average spectra of each clusters for one healthy mammary gland used for the training in negative ion mode. Related to Figure 3.

**Table S2.** Overview of all the percentages obtained for the three bacterial strains in each mice tumor or healthy mammary gland imaged in both ion mode.

## References

1. on behalf of the Swedish Digital Twin Consortium, Björnsson, B., Borrebaeck, C., Elander, N., Gasslander, T., Gawel, D.R., Gustafsson, M., Jörnsten, R., Lee, E.J., Li, X., et al. (2020). Digital twins to personalize medicine. Genome Med 12, 4. 10.1186/s13073-019-0701-3.

2. Hernandez-Boussard, T., Macklin, P., Greenspan, E.J., Gryshuk, A.L., Stahlberg, E., Syeda-Mahmood, T., and Shmulevich, I. (2021). Digital twins for predictive oncology will be a paradigm shift for precision cancer care. Nat Med 27, 2065–2066. 10.1038/s41591-021-01558-5.

3. Shafto, M. (2010). Modeling, Simulation, Information Technology & Processing RoadMap National Aeronautics and Space Administration.

4. Jiang, Y., Yin, S., Li, K., Luo, H., and Kaynak, O. (2021). Industrial applications of digital twins. Phil. Trans. R. Soc. A. 379, 20200360. 10.1098/rsta.2020.0360.

5. Wu, C., Lorenzo, G., Hormuth, D.A., Lima, E.A.B.F., Slavkova, K.P., DiCarlo, J.C., Virostko, J., Phillips, C.M., Patt, D., Chung, C., et al. (2022). Integrating mechanism-based modeling with biomedical imaging to build practical digital twins for clinical oncology. Biophysics Reviews 3, 021304. 10.1063/5.0086789.

6. Meijer, C., Uh, H.-W., and El Bouhaddani, S. (2023). Digital Twins in Healthcare: Methodological Challenges and Opportunities. JPM 13, 1522. 10.3390/jpm13101522.

7. Croatti, A., Gabellini, M., Montagna, S., and Ricci, A. (2020). On the Integration of Agents and Digital Twins in Healthcare. J Med Syst 44, 161. 10.1007/s10916-020-01623-5.

8. Shu, H., Liang, R., Li, Z., Goodridge, A., Zhang, X., Ding, H., Nagururu, N., Sahu, M., Creighton, F.X., Taylor, R.H., et al. (2023). Twin-S: a digital twin for skull base surgery. Int J CARS 18, 1077–1084. 10.1007/s11548-023-02863-9.

9. Rouhollahi, A., Willi, J.N., Haltmeier, S., Mehrtash, A., Straughan, R., Javadikasgari, H., Brown, J., Itoh, A., De La Cruz, K.I., Aikawa, E., et al. (2023). CardioVision: A fully automated deep learning package for medical image segmentation and reconstruction generating digital twins for patients with aortic stenosis. Computerized Medical Imaging and Graphics 109, 102289. 10.1016/j.compmedimag.2023.102289.

10. Servin, F., Collins, J.A., Heiselman, J.S., Frederick-Dyer, K.C., Planz, V.B., Geevarghese, S.K., Brown, D.B., Jarnagin, W.R., and Miga, M.I. (2024). Simulation of Image-Guided Microwave Ablation Therapy Using a Digital Twin Computational Model. IEEE Open J. Eng. Med. Biol. 5, 107–124. 10.1109/OJEMB.2023.3345733.

11. Tardini, E., Zhang, X., Canahuate, G., Wentzel, A., Mohamed, A.S.R., Van Dijk, L., Fuller, C.D., and Marai, G.E. (2021). Optimal policy determination in sequential systemic and locoregional therapy of oropharyngeal squamous carcinomas: A patient-physician digital twin dyad with deep Q-learning for treatment selection (Health Informatics) 10.1101/2021.04.07.21255092.

12. Rojas, I., Valenzuela, O., Rojas, F., Herrera, L.J., and Ortuño, F. eds. (2020). Bioinformatics and Biomedical Engineering: 8th International Work-Conference, IWBBIO 2020, Granada, Spain, May 6–8, 2020, Proceedings (Springer International Publishing) 10.1007/978-3-030-45385-5.

13. Bagaria, N. (2020). Health 4.0: Digital Twins for Health and Well-Being. Springer International Publishing.

14. Arai, K. ed. (2021). Intelligent Computing: Proceedings of the 2021 Computing Conference, Volume 3 (Springer International Publishing) 10.1007/978-3-030-80129-8.

15. Bruynseels, K., Santoni De Sio, F., and Van Den Hoven, J. (2018). Digital Twins in Health Care: Ethical Implications of an Emerging Engineering Paradigm. Front. Genet. 9, 31. 10.3389/fgene.2018.00031.

16. Corral-Acero, J., Margara, F., Marciniak, M., Rodero, C., Loncaric, F., Feng, Y., Gilbert, A., Fernandes, J.F., Bukhari, H.A., Wajdan, A., et al. (2020). The ‘Digital Twin’ to enable the vision of precision cardiology. European Heart Journal 41, 4556–4564. 10.1093/eurheartj/ehaa159.

17. Kaul, R., Ossai, C., Forkan, A.R.M., Jayaraman, P.P., Zelcer, J., Vaughan, S., and Wickramasinghe, N. (2023). The role of AI for developing digital twins in healthcare: The case of cancer care. WIREs Data Min & Knowl 13, e1480. 10.1002/widm.1480.

18. Vaysse, P.-M., Heeren, R.M.A., Porta, T., and Balluff, B. (2017). Mass spectrometry imaging for clinical research – latest developments, applications, and current limitations. Analyst 142, 2690–2712. 10.1039/C7AN00565B.

19. Duhamel, M., Drelich, L., Wisztorski, M., Aboulouard, S., Gimeno, J.-P., Ogrinc, N., Devos, P., Cardon, T., Weller, M., Escande, F., et al. (2022). Spatial analysis of the glioblastoma proteome reveals specific molecular signatures and markers of survival. Nat Commun 13, 6665. 10.1038/s41467-022-34208-6.

20. Buchberger, A.R., DeLaney, K., Johnson, J., and Li, L. (2018). Mass Spectrometry Imaging: A Review of Emerging Advancements and Future Insights. Anal. Chem. 90, 240–265. 10.1021/acs.analchem.7b04733.

21. Neumann, E.K., Djambazova, K.V., Caprioli, R.M., and Spraggins, J.M. (2020). Multimodal Imaging Mass Spectrometry: Next Generation Molecular Mapping in Biology and Medicine. J. Am. Soc. Mass Spectrom. 31, 2401–2415. 10.1021/jasms.0c00232.

22. Calligaris, D., Norton, I., Feldman, D.R., Ide, J.L., Dunn, I.F., Eberlin, L.S., Cooks, R.G., Jolesz, F.A., Golby, A.J., Santagata, S., et al. (2013). Mass Spectrometry Imaging as a Tool for Surgical Decision-Making. J Mass Spectrom 48, 1178–1187. 10.1002/jms.3295.

23. Tzafetas, M., Mitra, A., Paraskevaidi, M., Bodai, Z., Kalliala, I., Bowden, S., Lathouras, K., Rosini, F., Szasz, M., Savage, A., et al. (2020). The intelligent knife (iKnife) and its intraoperative diagnostic advantage for the treatment of cervical disease. PNAS 117, 7338– 7346. 10.1073/pnas.1916960117.

24. Saudemont, P., Quanico, J., Robin, Y.-M., Baud, A., Balog, J., Fatou, B., Tierny, D., Pascal, Q., Minier, K., Pottier, M., et al. (2018). Real-Time Molecular Diagnosis of Tumors Using Water-Assisted Laser Desorption/Ionization Mass Spectrometry Technology. Cancer Cell 34, 840–851.e4. 10.1016/j.ccell.2018.09.009.

25. Ifa, D.R., and Eberlin, L.S. (2016). Ambient Ionization Mass Spectrometry for Cancer Diagnosis and Surgical Margin Evaluation. Clinical Chemistry 62, 111–123. 10.1373/clinchem.2014.237172.

26. Ogrinc, N., Saudemont, P., Takats, Z., Salzet, M., and Fournier, I. (2021). Cancer Surgery 2.0: Guidance by Real-Time Molecular Technologies. Trends in Molecular Medicine 27, 602–615. 10.1016/j.molmed.2021.04.001.

27. Ogrinc, N., Saudemont, P., Balog, J., Robin, Y.-M., Gimeno, J.-P., Pascal, Q., Tierny, D., Takats, Z., Salzet, M., and Fournier, I. (2019). Water-assisted laser desorption/ionization mass spectrometry for minimally invasive in vivo and real-time surface analysis using SpiderMass. Nat Protoc 14, 3162–3182. 10.1038/s41596-019-0217-8.

28. Fatou, B., Saudemont, P., Leblanc, E., Vinatier, D., Mesdag, V., Wisztorski, M., Focsa, C., Salzet, M., Ziskind, M., and Fournier, I. (2016). In vivo Real-Time Mass Spectrometry for Guided Surgery Application. Sci Rep 6, 25919. 10.1038/srep25919.

29. Ledoux, L., Zirem, Y., Renaud, F., Duponchel, L., Salzet, M., Ogrinc, N., and Fournier, I. (2023). Comparing MS imaging of lipids by WALDI and MALDI: two technologies for evaluating a common ground truth in MS imaging. Analyst 148, 4982–4986. 10.1039/D3AN01096A.

30. Ogrinc, N., Kruszewski, A., Chaillou, P., Saudemont, P., Lagadec, C., Salzet, M., Duriez, C., and Fournier, I. (2021). Robot-Assisted SpiderMass for *In Vivo* Real-Time Topography Mass Spectrometry Imaging. Anal. Chem. 93, 14383–14391. 10.1021/acs.analchem.1c01692.

31. Balog, J., Szaniszlo, T., Schaefer, K.-C., Denes, J., Lopata, A., Godorhazy, L., Szalay, D., Balogh, L., Sasi-Szabo, L., Toth, M., et al. (2010). Identification of Biological Tissues by Rapid Evaporative Ionization Mass Spectrometry. Anal. Chem. 82, 7343–7350. 10.1021/ac101283x.

32. Eberlin, L.S., Norton, I., Orringer, D., Dunn, I.F., Liu, X., Ide, J.L., Jarmusch, A.K., Ligon, K.L., Jolesz, F.A., Golby, A.J., et al. (2013). Ambient mass spectrometry for the intraoperative molecular diagnosis of human brain tumors. Proc. Natl. Acad. Sci. U.S.A. 110, 1611–1616. 10.1073/pnas.1215687110.

33. Gredell, D.A., Schroeder, A.R., Belk, K.E., Broeckling, C.D., Heuberger, A.L., Kim, S.-Y., King, D.A., Shackelford, S.D., Sharp, J.L., Wheeler, T.L., et al. (2019). Comparison of Machine Learning Algorithms for Predictive Modeling of Beef Attributes Using Rapid Evaporative Ionization Mass Spectrometry (REIMS) Data. Sci Rep 9, 5721. 10.1038/s41598-019-40927-6.

34. Zirem, Y., Ledoux, L., Roussel, L., Maurage, C.A., Tirilly, P., Le Rhun, É., Meresse, B., Yagnik, G., Lim, M.J., Rothschild, K.J., et al. (2024). Real-time glioblastoma tumor microenvironment assessment by SpiderMass for improved patient management. Cell Reports Medicine, 101482. 10.1016/j.xcrm.2024.101482.

35. Nejman, D., Livyatan, I., Fuks, G., Gavert, N., Zwang, Y., Geller, L.T., Rotter-Maskowitz, A., Weiser, R., Mallel, G., Gigi, E., et al. (2020). The human tumor microbiome is composed of tumor type–specific intracellular bacteria. Science 368, 973–980. 10.1126/science.aay9189.

36. Galeano Niño, J.L., Wu, H., LaCourse, K.D., Kempchinsky, A.G., Baryiames, A., Barber, B., Futran, N., Houlton, J., Sather, C., Sicinska, E., et al. (2022). Effect of the intratumoral microbiota on spatial and cellular heterogeneity in cancer. Nature 611, 810–817. 10.1038/s41586-022-05435-0.

37. Long, Y., Tang, L., Zhou, Y., Zhao, S., and Zhu, H. (2023). Causal relationship between gut microbiota and cancers: a two-sample Mendelian randomisation study. BMC Med 21, 66. 10.1186/s12916-023-02761-6.

38. Xuan, C., Shamonki, J.M., Chung, A., DiNome, M.L., Chung, M., Sieling, P.A., and Lee, D.J. (2014). Microbial Dysbiosis Is Associated with Human Breast Cancer. PLoS ONE 9, e83744. 10.1371/journal.pone.0083744.

39. Dieleman, S., Aarnoutse, R., Ziemons, J., Kooreman, L., Boleij, A., and Smidt, M. (2021). Exploring the Potential of Breast Microbiota as Biomarker for Breast Cancer and Therapeutic Response. The American Journal of Pathology 191, 968–982. 10.1016/j.ajpath.2021.02.020.

40. Fernández, M., Reina-Pérez, I., Astorga, J., Rodríguez-Carrillo, A., Plaza-Díaz, J., and Fontana, L. (2018). Breast Cancer and Its Relationship with the Microbiota. IJERPH 15, 1747. 10.3390/ijerph15081747.

41. Alpuim Costa, D., Nobre, J.G., Batista, M.V., Ribeiro, C., Calle, C., Cortes, A., Marhold, M., Negreiros, I., Borralho, P., Brito, M., et al. (2021). Human Microbiota and Breast Cancer—Is There Any Relevant Link?—A Literature Review and New Horizons Toward Personalised Medicine. Front. Microbiol. 12, 584332. 10.3389/fmicb.2021.584332.

42. Green, J.E., Shibata, M.-A., Yoshidome, K., Liu, M., Jorcyk, C., Anver, M.R., Wigginton, J., Wiltrout, R., Shibata, E., Kaczmarczyk, S., et al. (2000). The C3(1)/SV40 T-antigen transgenic mouse model of mammary cancer: ductal epithelial cell targeting with multistage progression to carcinoma. Oncogene 19, 1020–1027. 10.1038/sj.onc.1203280.

43. Ogrinc, N., Saudemont, P., Balog, J., Robin, Y.-M., Gimeno, J.-P., Pascal, Q., Tierny, D., Takats, Z., Salzet, M., and Fournier, I. (2019). Water-assisted laser desorption/ionization mass spectrometry for minimally invasive in vivo and real-time surface analysis using SpiderMass. Nat Protoc 14, 3162–3182. 10.1038/s41596-019-0217-8.

44. Ruttkies, C., Schymanski, E.L., Wolf, S., Hollender, J., and Neumann, S. (2016). MetFrag relaunched: incorporating strategies beyond in silico fragmentation. J Cheminform 8, 3. 10.1186/s13321-016-0115-9.

45. Chambers, M.C., Maclean, B., Burke, R., Amodei, D., Ruderman, D.L., Neumann, S., Gatto, L., Fischer, B., Pratt, B., Egertson, J., et al. (2012). A cross-platform toolkit for mass spectrometry and proteomics. Nat Biotechnol 30, 918–920. 10.1038/nbt.2377.

46. Ke, G., Meng, Q., Finley, T., Wang, T., Chen, W., Ma, W., Ye, Q., and Liu, T.-Y. LightGBM: A Highly Efficient Gradient Boosting Decision Tree.

