## Supplemental figures legends and Tables for "Development of Molecular Digital Twins Based on Ambient Ionization Mass Spectrometry Imaging for Application in Cancer Surgery"

**Table S1. TgC(1)3 mice cohort.** Overview of the transgenic TgC(1)3 mice cohort available for the training and the blind testing of the classification model in both MS ionization modes. Related to Figures 2 and 3.

| **Mice number** | **Number of tumors per mice** | **Reference number** | **MS**  **Ionization mode** | **Use for training or blind analysis** |
| --- | --- | --- | --- | --- |
| TgC(1)3 N°1 | 1 | M1-T1 | Positive | Training |
| TgC(1)3 N°2 | 1 | M2-T1 | Positive | Training |
| TgC(1)3 N°3 | 3 | M3-T1, M3-T2, M3-T3 | Negative | Training |
| TgC(1)3 N°4 | 1 | M4-T1 | Negative | Training |
| TgC(1)3 N°7 | 2 | M7-T1, M7-T2 | Negative | Training |
| TgC(1)3 N°8 | 2 | M8-T1, M8-T2 | Negative | Training |
| TgC(1)3 N°9 | 2 | M9-T1, M9-T2 | Positive | Training and test in blind |
| TgC(1)3 N°11 | 1 | M11-T1 | Negative | Test in blind |
| TgC(1)3 N°12 | 1 | M12-T1 | Negative | Training |
| TgC(1)3 N°13 | 1 | M13-T1 | Negative | Training |
| TgC(1)3 N°14 | 2 | M14-T1, M14-T2 | Negative | Training |
| TgC(1)3 N°15 | 2 | M15-T1, M15-T2 | Negative | Training and test in blind |
| Healthy | 1 | Healthy | Negative | Training |

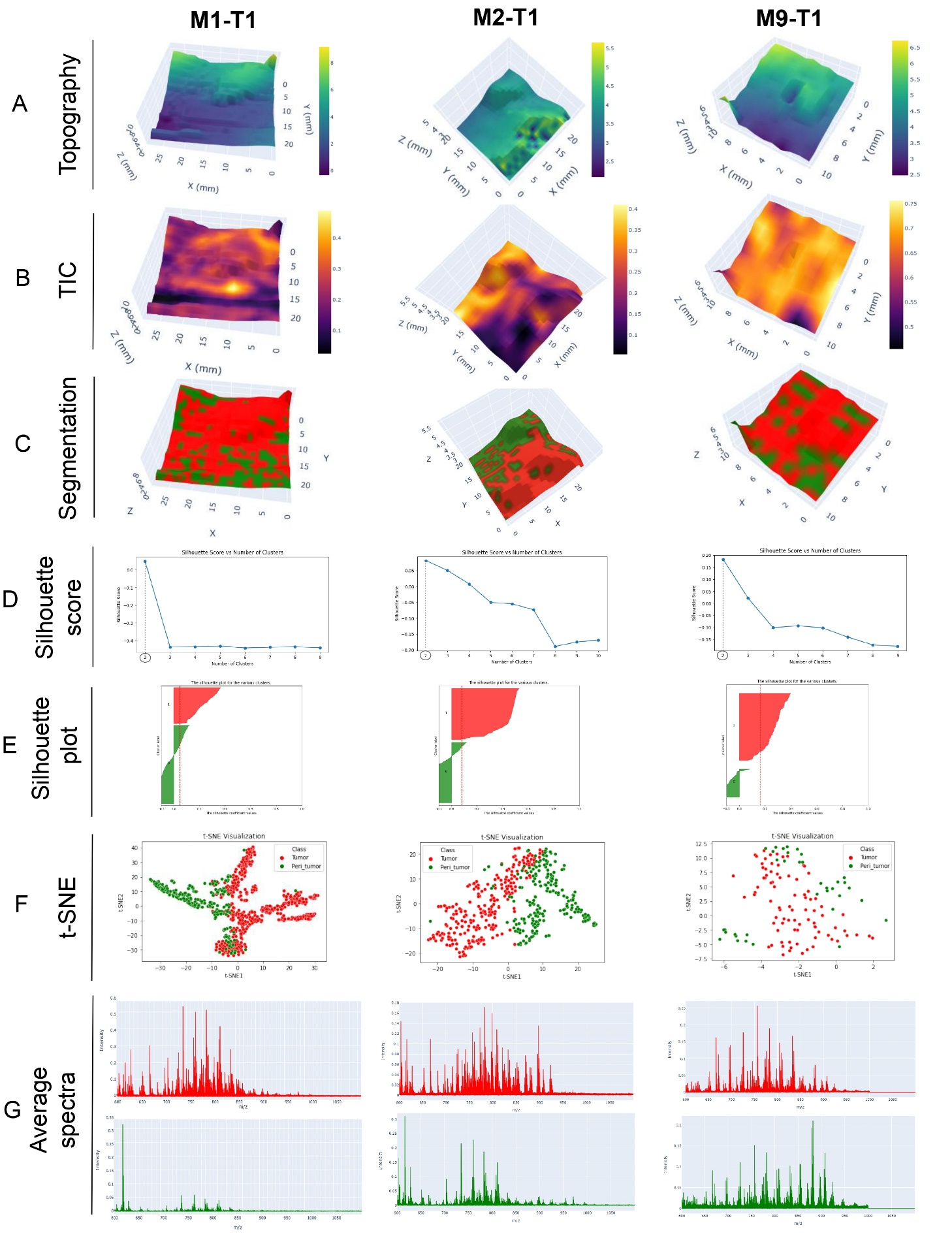

**Figure S1. Topographic MSI data in the positive ion mode of M1-T1, M2-T1 and M9-T1 tumors.** (A) Topographic map, (B) TIC map, (C) segmentation map, (D) silhouette score, (E) silhouette plot, (F) t-SNE and (G) average spectra of each cluster for 3 mice tumors (M1-T1, M2-T1 and M9-T1) used for the training in positive ion mode. Related to Figure 2.

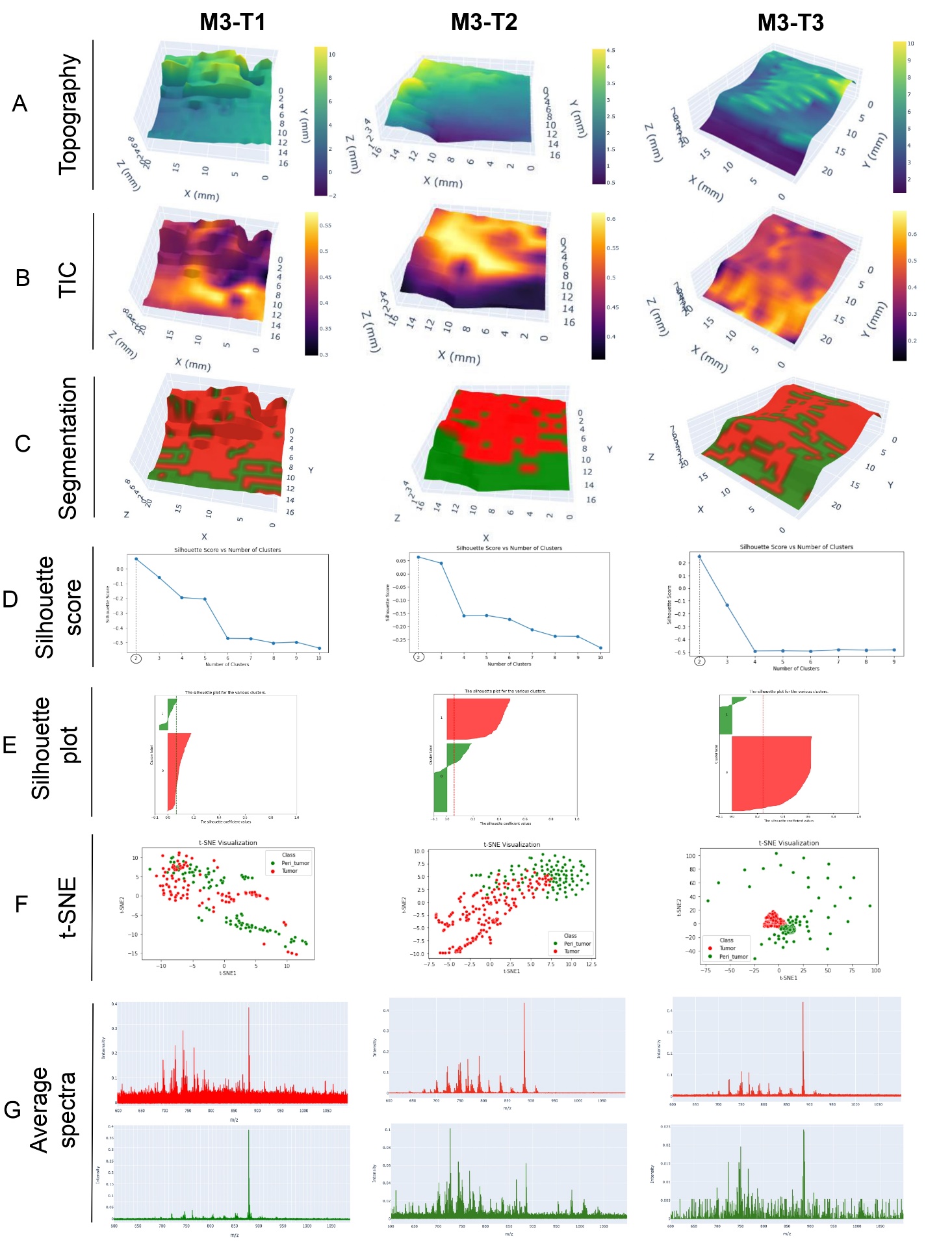

**Figure S2.** **Topographic MSI data in the negative ion mode of M3-T1, M3-T2 and M3-T3 tumors.** (A) Topographic map, (B) TIC map, (C) segmentation map, (D) silhouette score, (E) silhouette plot, (F) t-SNE and (G) average spectra of each cluster 3 mice tumors (M3-T1, M3-T2 and M3-T3) used for the training in negative ion mode. Related to Figure 3.

.
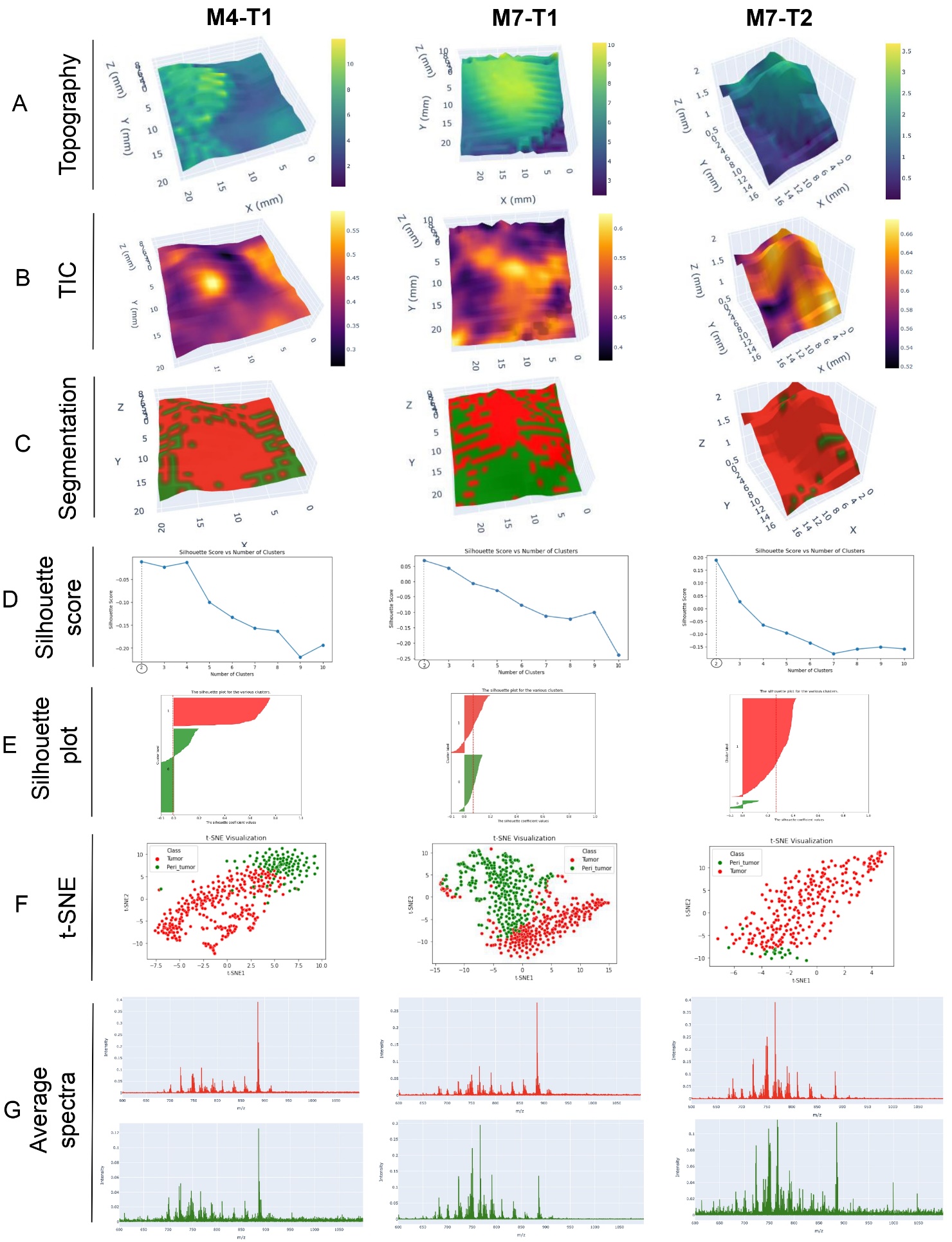

**Figure S3.** **Topographic MSI data in the negative ion mode of M4-T1, M7-T1 and M7-T2 tumors.** (A) Topographic map, (B) TIC map, (C) segmentation map, (D) silhouette score, (E) silhouette plot, (F) t-SNE and (G) average spectra of each cluster 3 mice tumors (M4-T1, M7-T1 and M7-T2) used for the training in negative ion mode. Related to Figure 3

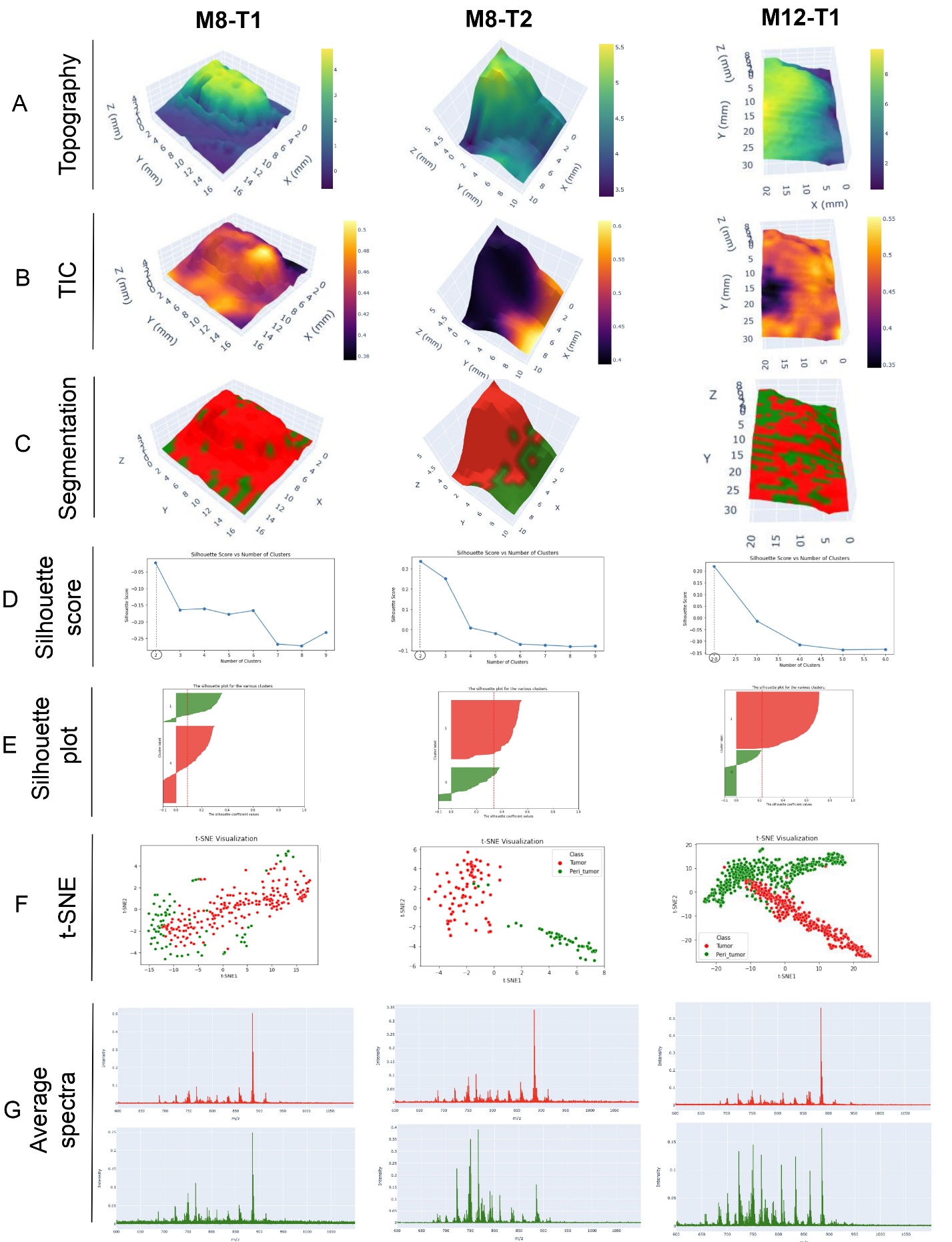

**Figure S4.** **Topographic MSI data in the negative ion mode of M8-T1, M8-T2 and M2-T1 tumors.** (A) Topographic map, (B) TIC map, (C) segmentation map, (D) silhouette score, (E) silhouette plot, (F) t-SNE and (G) average spectra of each cluster 3 mice tumors (M8-T1, M8-T2 and M12-T1) used for the training in negative ion mode. Related to Figure 3.

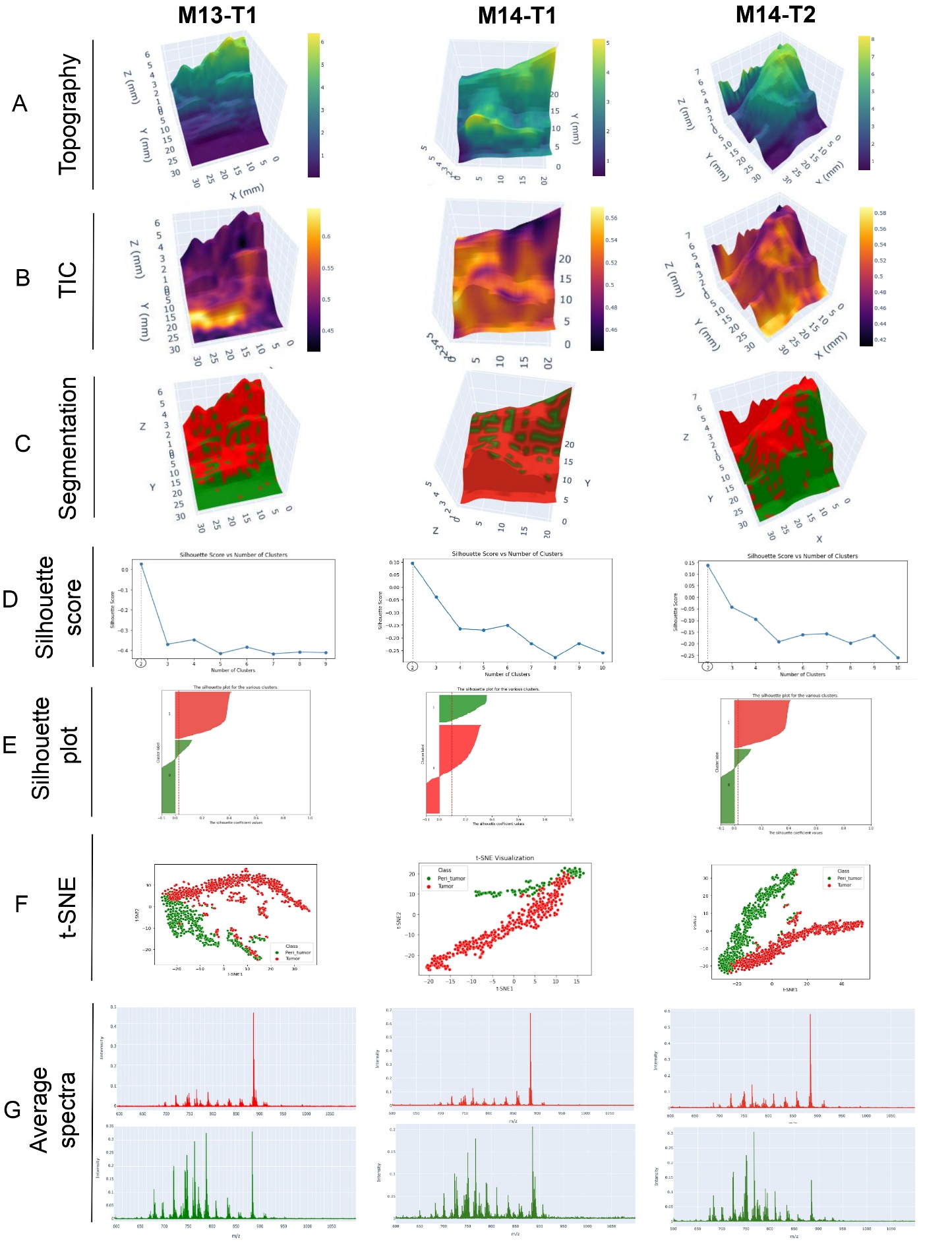

**Figure S5.** **Topographic MSI data in the negative ion mode of M13-T1, M14-T1 and M14-T2 tumors.** (A) Topographic map, (B) TIC map, (C) segmentation map, (D) silhouette score, (E) silhouette plot, (F) t-SNE and (G) average spectra of each cluster 3 mice tumors (M13-T1, M14-T1 and M14-T2) used for the training in negative ion mode. Related to Figure 3.

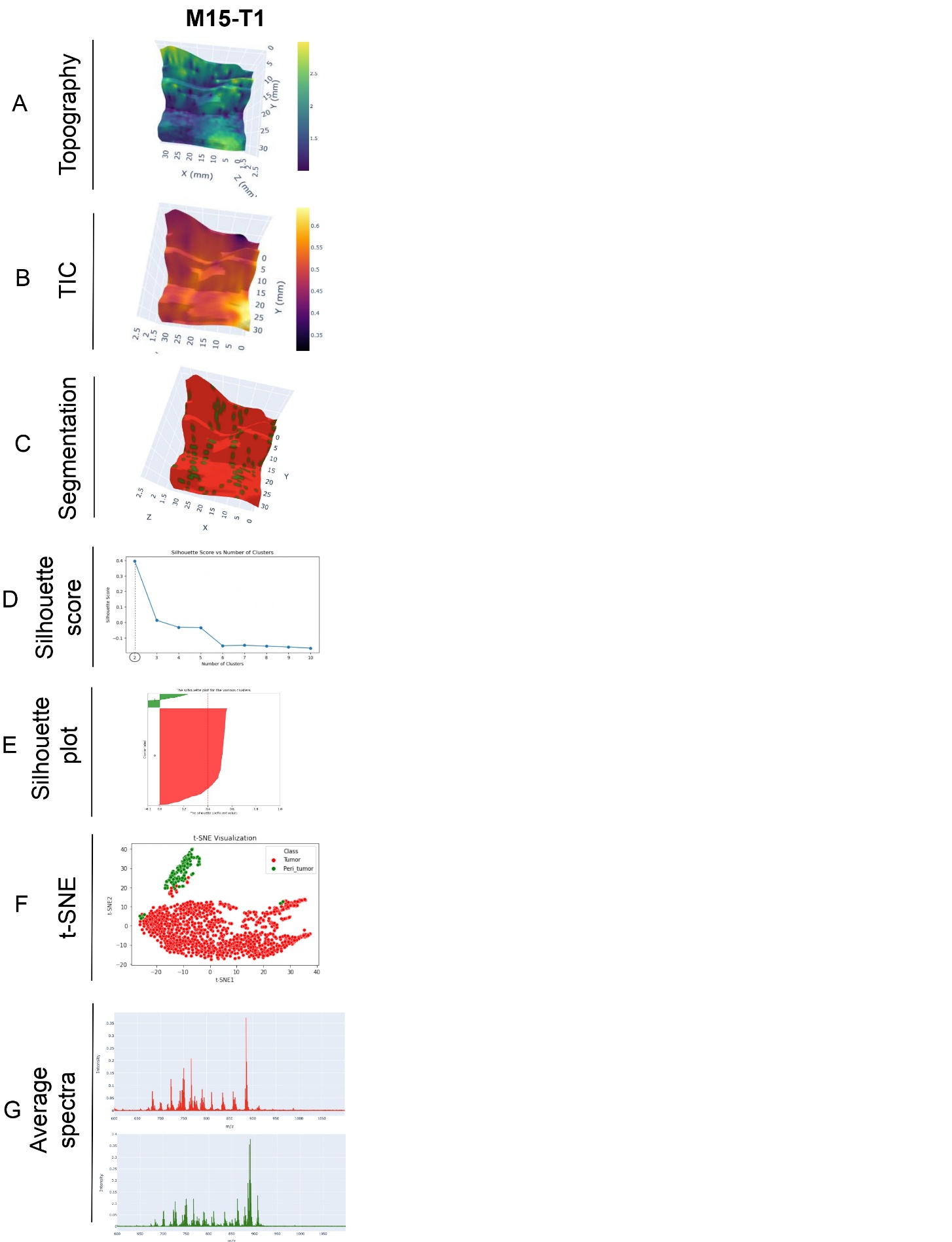

**Figure S6.** **Topographic MSI data in the negative ion mode of M5-T1 tumor.** (A) Topographic map, (B) TIC map, (C) segmentation map, (D) silhouette score, (E) silhouette plot, (F) t-SNE and (G) average spectra of each cluster 1 mice tumor (M15-T1) used for the training in negative ion mode. Related to Figure 3.

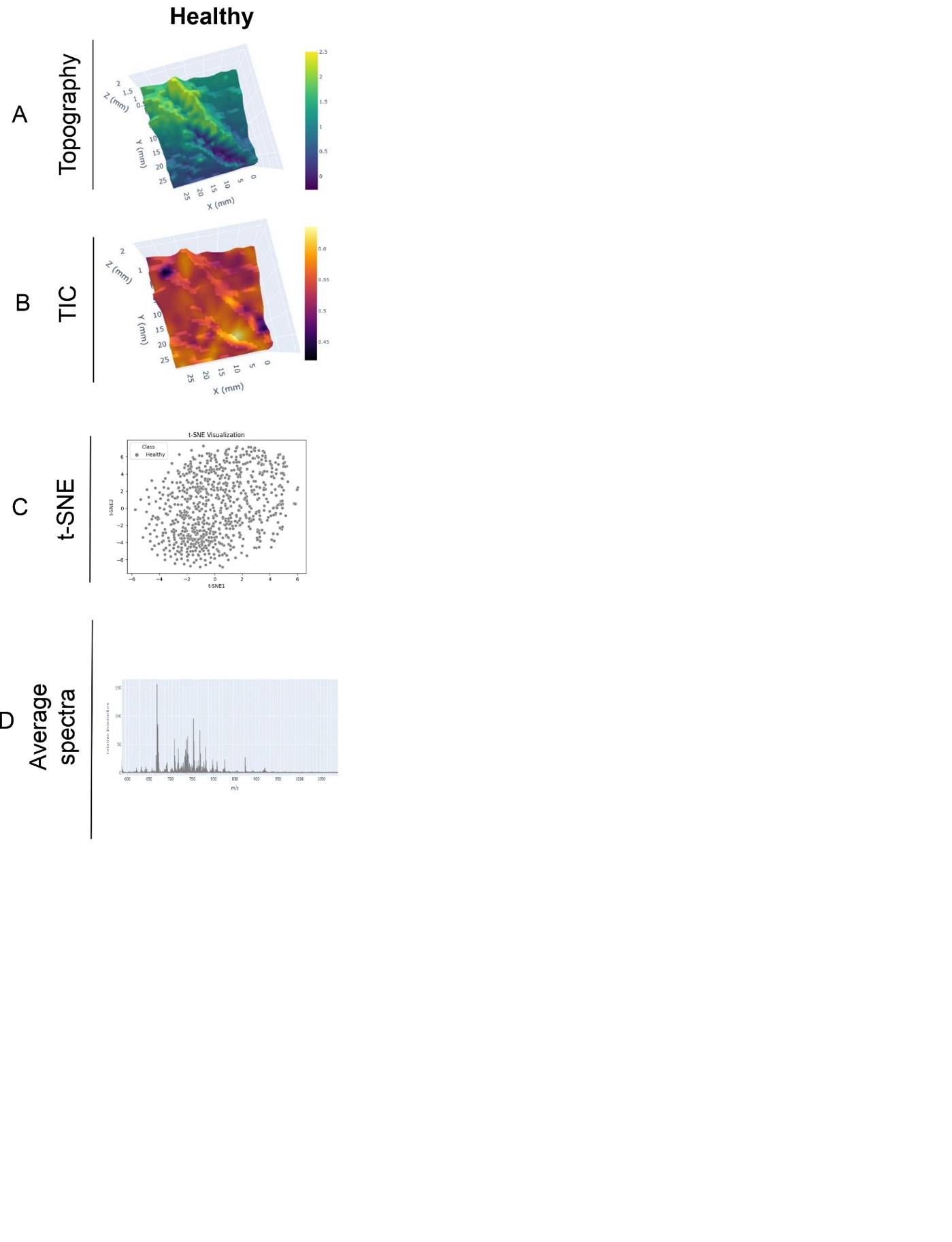

**Figure S7.** **Topographic MSI data in the negative ion mode of a healthy mammary.** (A) Topographic map, (B) TIC map, (C) segmentation map, (D) silhouette score, (E) silhouette plot, (F) t-SNE and (G) average spectra of each cluster for one healthy mammary gland used for the training in negative ion mode. Related to Figure 3.

**Table S2. Percentages of *S. infantis, S. lugdunensis, M. radiotolerans bacteria* in the tumor and peritumoral region of the different mice tumors.** Percentages obtained for the three bacterial strains in each mammary gland tissues imaged by SpiderMass in both MS ion modes. Related to Figure 6.

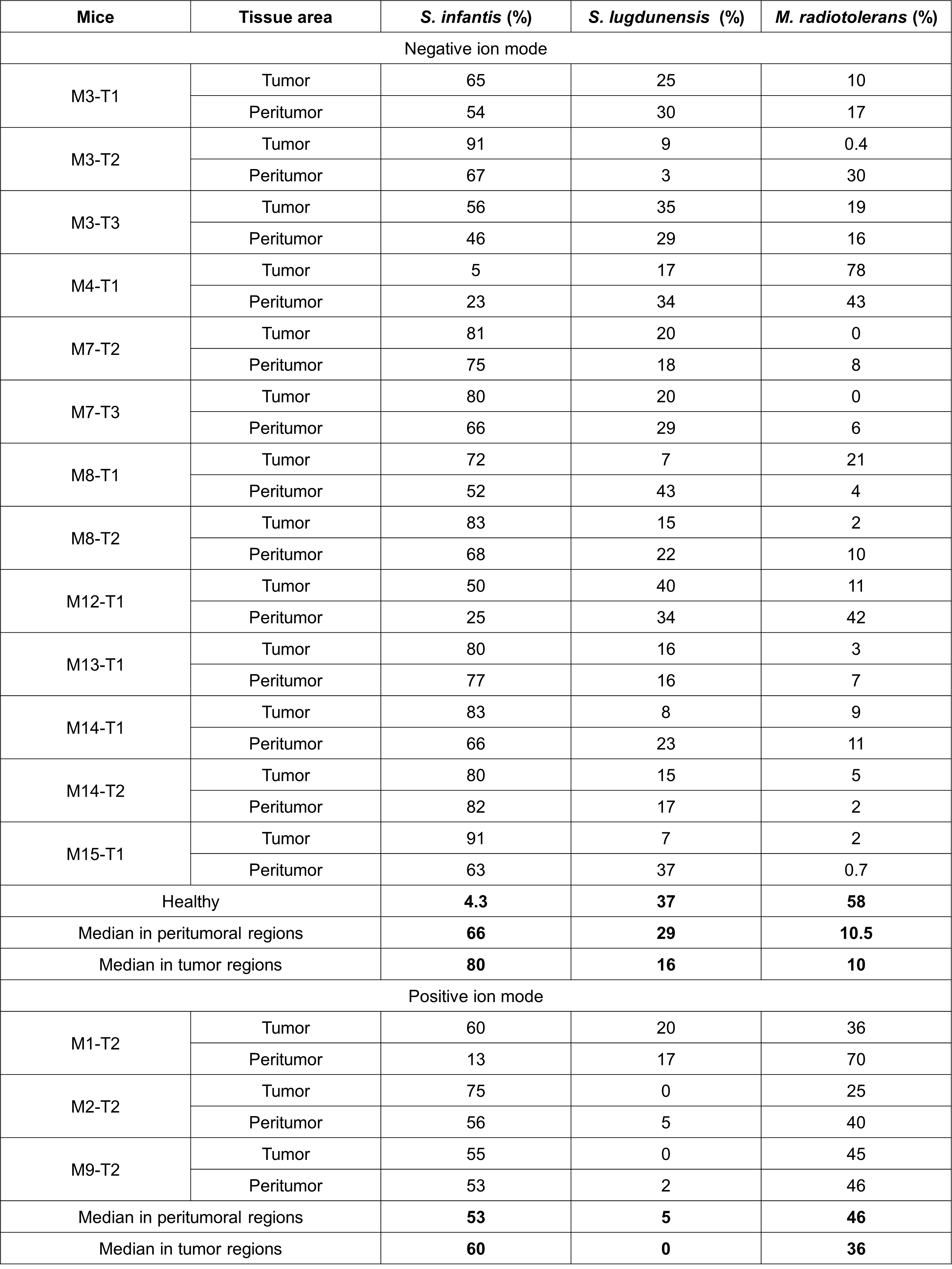

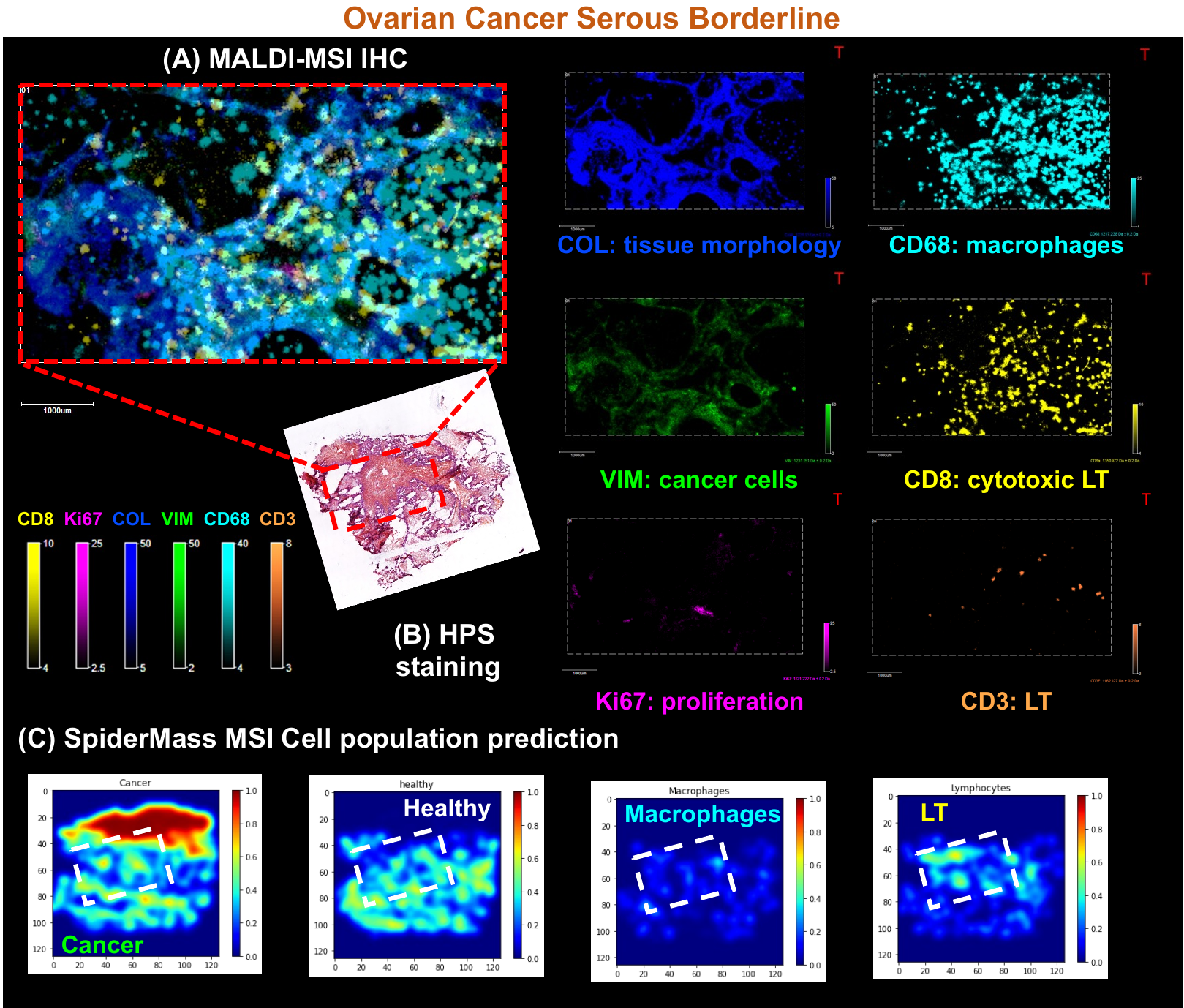

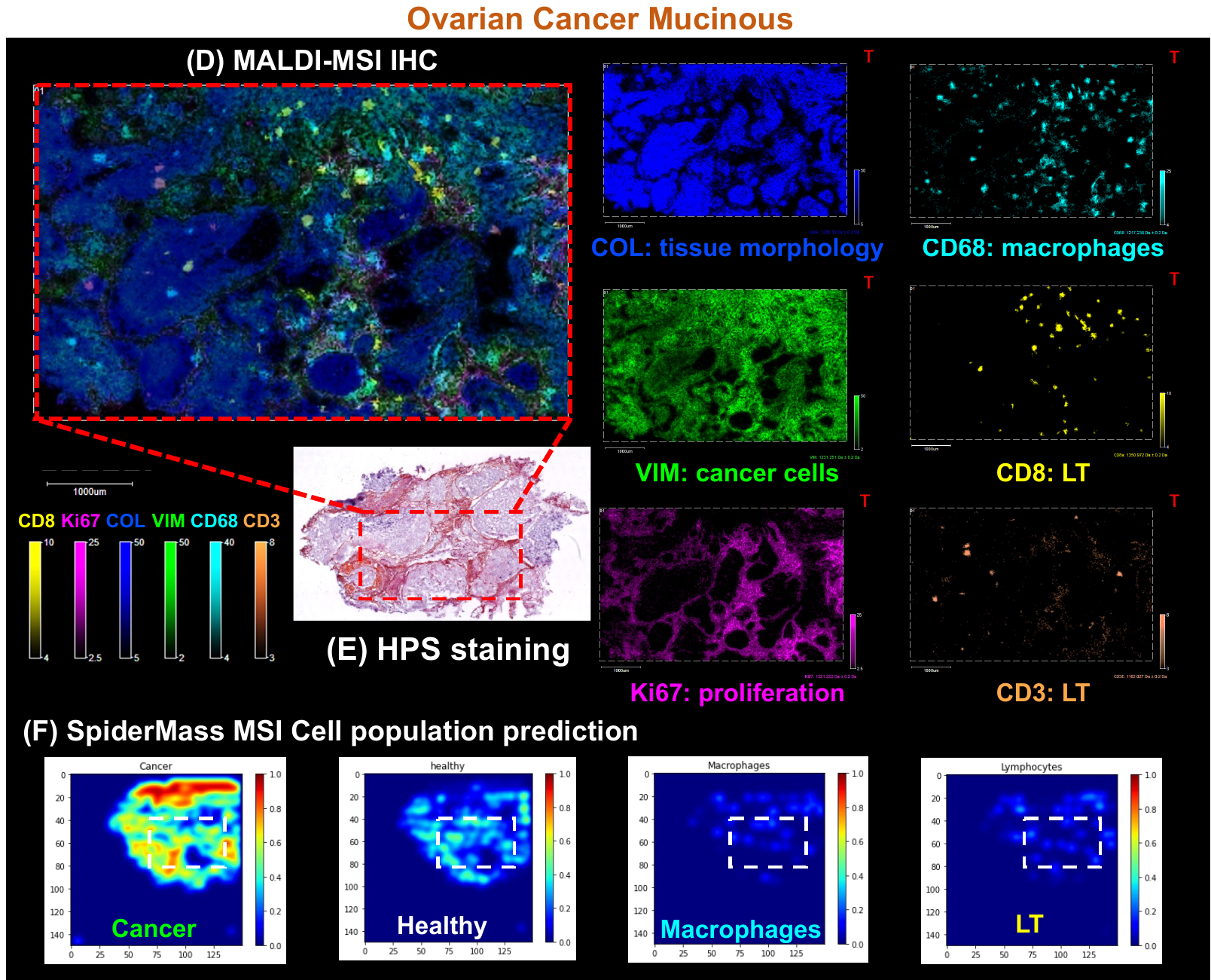

**Figure S8.** **Cross-validation of the different cell population predictions from the SpiderMass MSI data by MALDI-MSI IHC against markers of normal, cancer and immune cells.** MALDI MSI-IHC (A, C) in 6-plex against CD8 (Lymphocytes T cytotoxic), Ki67 (proliferation), collagen (tissue morphology), vimentin (cancer cells), CD68 (macrophages), CD3 (Lymphocyte T); HPS staining (B, E) and different cell populations prediction (cancer, normal cells, macrophages, LT) based on SpiderMass MSI data (C, F). Related to Figure 5.
